# The composition and function of *Enterococcus faecalis* membrane vesicles

**DOI:** 10.1101/2021.01.28.428366

**Authors:** Irina Afonina, Brenda Tien, Zeus Nair, Artur Matysik, Ling Ning Lam, Mark Veleba, Augustine Koh, Rafi Rashid, Amaury Cazenave-Gassiot, Marcus Wenk, Sun Nyunt Wai, Kimberly A. Kline

## Abstract

Membrane vesicles (MVs) contribute to various biological processes in bacteria, including virulence factor delivery, host immune evasion, and cross-species communication. MVs are frequently being discharged from the surface of both Gram-negative and Gram-positive bacteria during growth. In some Gram-positive bacteria, genes affecting MV biogenesis have been identified, but the mechanism of MV formation is unknown. In *Enterococcus faecalis*, a causative agent of life-threatening bacteraemia and endocarditis, neither mechanisms of MV formation nor their role in virulence has been examined. Since MVs of many bacterial species are implicated in host-pathogen interactions, biofilm formation, horizontal gene transfer, and virulence factor secretion in other species, we sought to identify, describe, and functionally characterize MVs from *E. faecalis*. Here we show that *E. faecalis* releases MVs that possess unique lipid and protein profiles, distinct from the intact cell membrane, and are enriched in lipoproteins. MVs of *E. faecalis* are specifically enriched in unsaturated lipids that might provide membrane flexibility to enable MV formation, providing the first insights into the mechanism of MV formation in this Gram-positive organism.

## Introduction

Membrane vesicles (MVs) are widely produced by Gram-negative and Gram-positive bacteria. While MVs confer many similar functions to Gram-positive and Gram-negative bacteria, including virulence factor delivery and host immune modulation, the mechanism of MV formation differs due to intrinsic differences in the structure of the cell wall [1–5]. MVs are well-studied in Gram-negative bacteria where they are derived from the outer membrane and are produced under the normal growth and stress conditions [6–9]. Since Gram-positive bacteria lack an outer membrane and have a thicker peptidoglycan which could impact MV formation or release, MVs in Gram-positive bacteria were largely neglected until the first report in 2009 characterizing MVs from *Staphylococcus aureus* [10]. Since then, MVs have been reported in numerous Gram-positive bacteria including *Bacillus subtilis*, *Streptococcus pyogenes*, *Streptococcus mutans*, *Listeria monocytogenes*, *Enterococcus faecium,* but not in *Enterococcus faecalis* [3, 10–14]. While several mechanisms for outer membrane vesicle (OMV) formation in Gram-negative bacteria have been proposed, the mechanism of MV formation in Gram-positive bacteria is largely unknown [15]. In Gram-positive bacteria, MV biogenesis can be regulated genetically. In *L. monocytogenes*, RNA polymerase sigma factor σB, a general stress transcription factor that activates stress response genes and *ftsZ*, contributes to MV formation with significantly reduced vesiculation in a Δ*sigB* mutant [14]. Similarly, MV biogenesis in *M. tuberculosis* is affected by VirA (vesiculogenesis and immune response regulator) and vesiculation increases in a VirA deficient mutant [16]. In *S. pyogenes,* the two-component system CovRS negatively impacts vesiculation [17].

*E. faecalis* is an opportunistic pathogen that can cause urinary tract infections (UTI), catheter associated urinary tract infections (CAUTI), wound infection, and life-threatening bacteraemia and endocarditis [18–20]. Infections by this opportunistic pathogen can be difficult to treat due to their propensity to form biofilms, as well as their frequent and multiple antibiotic resistances [21]. *E. faecalis* and *E. faecium* both cause disease in humans, but *E. faecalis* is more frequently isolated from clinical specimens [20]. Understanding the mechanism of vesiculation in *E. faecalis* and the role MVs play in virulence, may improve strategies to treat enterococcal infections.

## Methods

### Bacterial strains and growth conditions

*Enterococcus faecalis* OG1RF and *Escherichia coli* DH5α with or without the plasmid pGCP123 [22] were used in this study. *E. faecalis* strains were grown statically in brain heart infusion media (BHI; BD Difco, USA) or on BHI agar (BHI supplemented with 1.5% agarose (1st BASE, Singapore)) at 37°C. *E. coli* was grown in Luria-Bertani Broth (Miller) (LB; BD, Difco, USA) at 37°C, 200 rpm shaking. For biofilm assays, tryptone soy broth (Oxoid, UK) supplemented with 10 mM glucose (TSBG) was used. Kanamycin (kan) was used in the following concentrations where appropriate: 50 μg/mL (*E. coli*), 500 μg/mL (*E. faecalis*), unless otherwise noted.

### Genetic manipulations

To construct the in-frame deletion of pp2 (OG1RF 11046-11063), regions approximately 1Kb upstream and downstream of the genes were amplified from OG1RF using primer pairs pp2.infu.1F/R and pp2.infu.2F/R for upstream and downstream regions, respectively (**Table S1**). These products were sewn together and amplified using pp2.infu.1F / pp2.infu.2R. The temperature sensitive plasmid pGCP213 was amplified using the pGCPinphuF/R primer pair. The pp2.infuF/R PCR product of approximately 2Kb was then cloned into the pGCP213 fragment using the In-Fusion HD Cloning Plus System (Takara Bio Inc, Shimogyō-ku, Kyoto) to generate the temperature sensitive deletion plasmid pGCPpp2.

Deletion constructs were then transformed into OG1RF by electroporation and the transformants were selected at 30°C on agar plates with kan. Chromosomal integrants were selected by growth at 42°C on the agar plate in the presence of kan. Selection for excision of the integrated plasmid by homologous recombination was accomplished by growing the bacteria at 30°C in the absence of kan in the broth. Loss of the pp2 locus in kanamycin-sensitive bacteria was demonstrated by PCR using primer pair infu_check_pp2F/R

### Membrane vesicle isolation

A single *E. faecalis* colony was inoculated into 100 mL of BHI broth and grown overnight. The resulting overnight culture was diluted 1:10 with warm fresh BHI and grown for 2 hours. Cells were harvested using a Beckman centrifuge and JA 12.650 rotor at 4000 x g for 15 min at 4°C. Supernatants were filtered through a 0.45 μm vacuum filter and concentrated to 70 mL using a VIVAFLOW 100,000 MWCO Hydrosart (Sartorius, Germany). Concentrated supernatants were then subjected to ultracentrifugation at 160,000 x g for 2 hours at 4°C using a Beckman Ti 45 rotor (Beckman, Germany). Supernatants were completely removed and the pellet containing the crude MV fraction was resuspended in 400 μL of chilled PBS. MVs were further purified using density gradient centrifugation as previously described [23]. Briefly, 400 μL of PBS containing crude MVs was layered over an OptiPrep density gradient in a 4.2 mL tube in the following order from bottom to top: 400 μL (45% OptiPrep), 500 μL (35% OptiPrep), 600 μL (30% OptiPrep), 600 μL (25% OptiPrep), 600 μL (20% OptiPrep), 500 μL (15% OptiPrep), 600 μL (10% OptiPrep). The 4.2 mL tubes were centrifuged in a SW60 Ti rotor (Beckman Coulter, USA) for 3 hours at 160,000 x g at 4°C. After centrifugation, 200 μL OptiPrep gradient aliquots were removed from the top and transferred into 21 Eppendorf tubes (21 fractions in total). Collected fractions were subjected to SDS-PAGE and silver staining. Fractions 13-16 containing MVs were combined and used within two days or stored at – 80°C until further analysis. For Nanosight, proteomics, and lipidomic analyses, samples were further purified from OptiPrep by ultracentrifugation at 160,000 x g for 2 hours and the pellet was resuspended in 50 μL of PBS.

### Transmission electron microscopy (TEM) analysis

For negative staining, 4 μL of sample was deposited onto glow-discharged carbon-coated copper grids (NTU, Singapore). After 1 min, the sample was blotted with a filter paper and 4 μL of 1% uranyl acetate was added onto the grid. After 1 min, excess uranyl acetate was removed. Samples were visualized using a T12 120kV TEM (Fei, USA).

### Live imaging

Late-log *E. faecalis* sub-culture in BHI was washed in PBS, stained with Nile Red (10μg/mL) for 10 mins at room temperature, washed again in PBS, spotted on 1mm thick BHI agar (1.5%) pad prepared on a glass slide and mounted with coverslip (glass-agar-glass). Sample was then imaged using Zeiss Axio Observer 7 widefield epifluorescence microscope, equipped with 100x/1.4NA PlanApochromat oil-immersion objective and Hamamatsu Orca Flash 4.0 detector. Images were acquired with 2 seconds interval with minimal excitation light intensity but sufficient for imaging, to minimize photobleaching and phototoxicity effects. Images were then processed using ImageJ/FIJI.

### Nanosight analysis

Purified MV samples were analysed with a NanoSight NS300 (Malvern Instruments, UK) according to the manufacturer’s protocol. Briefly, the system was first washed 3 times with 1 mL sterile water. Next, the sample was diluted 1:10 in 1X PBS and loaded into the chamber with a sterile syringe to ensure no bubbles were trapped in the installed tubing. All samples were captured under the same settings and concentration and size were calculated automatically by NanoSight NS300 Control Software (Malvern Instruments, UK).

### MV plasmid transfer assay

250 μL of purified MV sample was added to 10^4^ CFU/mL of recipient OG1RF or DH5α in 250 μL of 2X BHI (for *E. faecalis* OG1RF) or 2X LB (for *E. coli* DH5α) broth. Two hours after incubation, 100 μL of recipient *E. faecalis* or *E. coli* DH5α was plated on BHI agar (kan 1000 μg/mL) for *E. faecalis* or LB agar *E. coli* (kan 50 μg/mL) in duplicates. The remaining volume of recipient *E. faecalis* OG1RF or *E. coli* DH5α was serially diluted plated on plain BHI/ LB agar, incubated overnight for CFU enumeration.

### DNA quantification

DNA concentration in MV samples before and after heating at 95°C for 10 mins was measured by Qubit (Qubit^®^ 2.0 Fluorometer, Invitrogen™, Thermo Fisher Scientific) as per user manufacturer’s instructions.

### DNAse treatment

Five μL of crude or purified MV sample was treated with 1 μL DNAse in a 50 μL reaction volume for 30 min at 37°C. 10 ng of purified pGCP123 was added to control samples. Where indicated, DNase was inactivated at 98°C for 10 min prior to addition of the sample and PCR reaction using primers to amplify pGCP123 plasmid: M13 Forward (5’-GTAAAACGACGGCCAGTG-3’)/ Reverse (5’-CAGGAAACAGCTATGAC-3’).

### Cell culture and NF-κB reporter assay

RAW-Blue cells derived from RAW 264.7 macrophages (Invivogen), containing a plasmid encoding a secreted embryonic alkaline phosphatase (SEAP) reporter under transcriptional control of an NF-κB-inducible promoter, were cultivated in Dulbecco Modified Eagle medium containing 4500 mg/L high glucose (1X) with 4.0 nM L-glutamine without sodium pyruvate (Gibco), and supplemented with 10% fetal bovine serum (FBS) (Gibco) supplemented with 200 μg/mL Zeocin at 37°C in 5% CO_2_.

RAW-Blue cells were seeded in a 96 well plate at 100,000 cells/well in 200 μL of antibiotic-free cell culture media. Following overnight incubation, the cells were washed once with PBS and fresh media was added. Cells were stimulated using LPS purified from *E. coli* O111:B4 (Sigma Aldrich) (100 ng/mL) as a positive control, or cell culture media alone or OptiPrep alone as negative controls. MVs purified in OptiPrep (1000 particles/macrophage) were added to RAW-Blue cells and incubated for 6 hours with or without simultaneous LPS stimulation. Post-infection, 20 μL of supernatant was added to 180 μL of QUANTI-Blue reagent (Invivogen) and incubated overnight at 37°C. SEAP levels were determined at 640 nm using a TECAN M200 microplate reader.

### Protein content analysis

Purified MV samples or mid-log phase intact bacterial cells, washed in PBS and incubated in lysis buffer with 1 mg/mL lysozyme for 1 hour at 37°C, were boiled in 1X NuPAGE^®^ LDS sample buffer (Invitrogen, USA) and 0.1 M dithiothreitol (DTT) at 95°C for 10 min. Samples were run on 10% NuPAGE^®^ Bis-Tris mini gel (ThermoFisher Scientific, USA) in 1x MOPS SDS running buffer in XCellSureLock^®^ Mini-Cell for 10 min at 150 V. Gel lanes containing silver stained proteins were cut out and sent to the Harvard Medical School Taplin Mass Spectrometry facility for mass spectrometry (MS) analysis (Harvard, USA). From the list of identified proteins within each sample, proteins containing fewer than 3 unique peptides were excluded from the analysis. All the other proteins were assigned an abundance score (A*) based on the number of unique peptides per protein divided by the total number of unique peptides. We averaged A* within triplicates and sorted proteins in descending order based on the A*. The most abundant proteins are the top 10 proteins based on the A*, the enriched proteins were identified as A* of the protein within MV sample divided by A* of the same protein within whole cell lysate fraction. If the protein was not present within the whole cell lysate fraction, A* was defined as 0.06, the lowest abundance within all proteins in WCL, when calculating enrichment.

### Lipid content analysis

Lipids were extracted from lyophilised membrane vesicles or whole cell pellets, using a modified Bligh & Dyer method as previously described [24, 25]. Prior to extraction, samples were spiked with known amounts of internal standards for phosphatidylglycerol (PG) and lysyl-phosphatidylglycerol (Lys-PG) or external calibration standard, monoglucosyl-diacylglycerol (MGDAG 34:1) (Avanti polar lipids, Alabaster, AL, USA) and were run alongside the samples (**Table S2**).

PG and L-PG in WCL and MVs were quantified by LC-MS/MS using multiple reaction monitoring (MRM) that we previously established [24]. An Agilent 6490 or 6495A QqQ mass spectrometer connected to a 1290 series chromatographic system was used together with electrospray ionization (ESI) for lipid ionisation. Each lipid molecular species was analyzed using a targeted MRM approach containing transitions for known precursor/product mass-to-charge ratio (m1/m3). Signal intensities were normalized to the spiked internal standards (PG 14:0 and L-PG 16:0) to obtain relative measurements, as described previously (13). Due to an absence of suitable internal standards, semi-quantitative analysis of diglucosyl-diacylglycerol (DGDAG) was carried out instead. Lipid extraction was performed without spiking of internal standards and DGDAG lipid species were analysed by LC-MS/MS using MRMs using monoglucosyl-diacylglycerol (MGDAG) 34:1 as a surrogate standard (**Table S2**) for external calibration curves. Measurements of MGDAG 34:1 dilution, from 0.2 ng/mL to 1000 ng/mL were used to construct external calibration curves to estimate the levels of DGDAG. The MRM transitions for DGDAG molecular species and MGDAG 34:1 are listed in **Table S3**. All lipid species abundances were expressed as a percentage of their respective lipid class.

### Statistical Analysis

Statistical analyses were performed using GraphPad Prism software (Version 6.05 for Windows, California, United States). All experiments were performed at least in three biological replicates and the mean value was calculated. All graphs indicate standard deviation from independent experiments. Statistical analysis was performed by unpaired t-test using GraphPad (* p<0.05, ** p< 0.01, *** p<0.001; **** p<0.0001, ns: p>0.05). SEAP assays were analyzed using one-way ANOVA with Tukey’s multiple comparison. *P*-values less than 0.05 were deemed significant.

## Results

### *E. faecalis* produces MVs with a size range of 50-400nm

To determine whether *E. faecalis* produces MVs, we used methods previously described for the isolation of MVs from Gram-positive bacteria [10]. We collected cell-free supernatants of late-log phase cultures and performed ultracentrifugation to obtain a crude pellet. We then separated the crude pellet by OptiPrep gradient centrifugation, from which we collected 22 fractions followed by SDS PAGE and silver stain (**Figure 1A**). We observed a denser staining pattern with altered protein abundances in fractions 13-16, at approximately 25% OptiPrep, a similar density at which MVs were found for *S. aureus* and *E. coli* [10, 26, 27].

**Figure 1.**
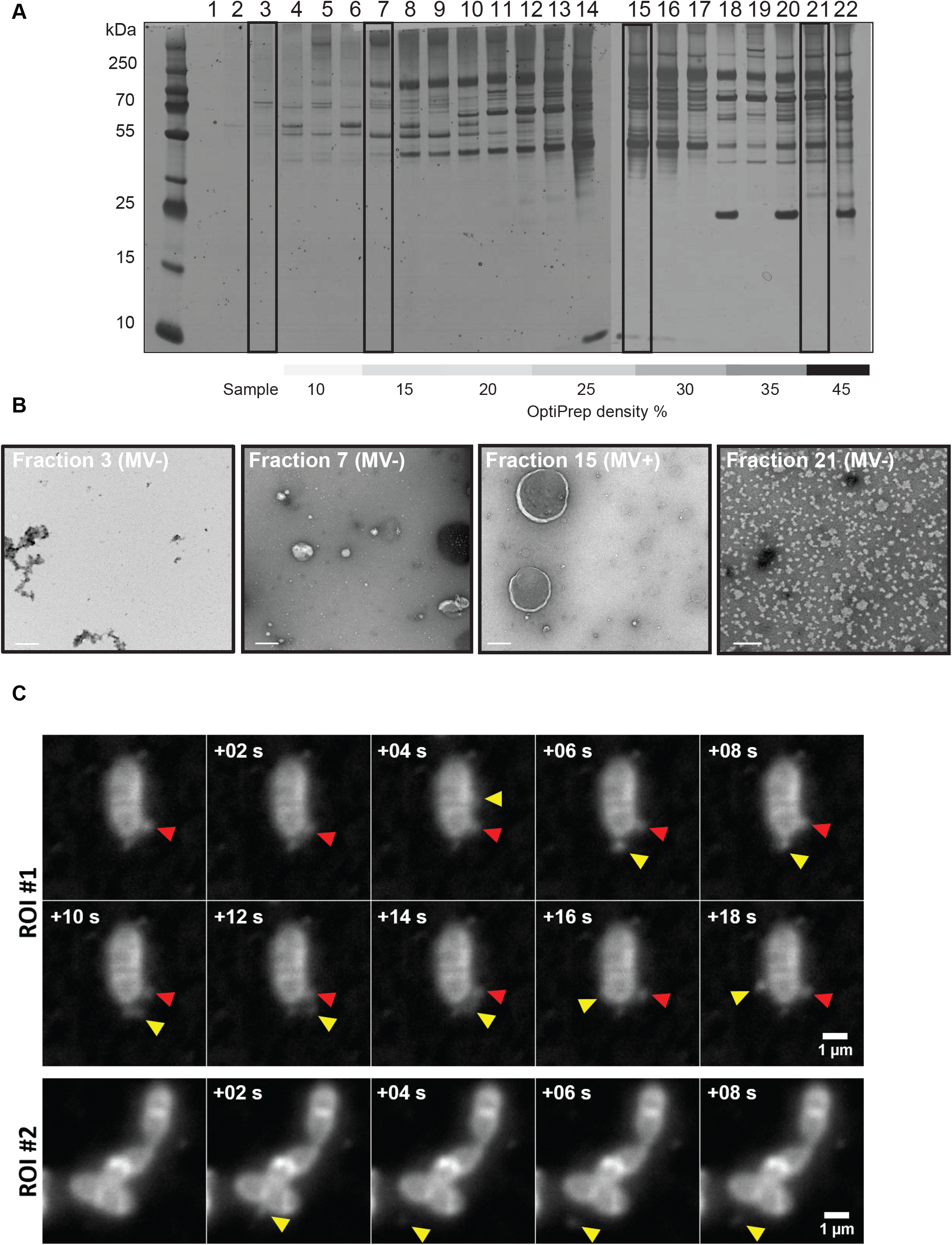
*E. faecalis* produces MVs ranging from 40-400 nm in size. (**A**) 22 fractions consisting of 200 μL each were collected from the top of a 4.4 mL OptiPrep gradient, separated by SDS-PAGE, and silver stained. (**B**) Selected fractions (# 3, 7, 15, 21 indicated in the boxes in panel A) were negative stained and viewed by TEM. Scale bar is 200 nm. (**C**) In-situ imaged live *E. faecalis* on the agar pad. Panels of consecutive frames (+0.2 sec) from two distinct area on the pad. Existing vesicle is indicated with a red arrow, the new vesicle - with a yellow arrow. Scale bar 1 μL.

To determine whether fractions 13-16 contained MVs, we performed transmission electron microscopy (TEM) on fraction 15 (MV fraction) as well as 3 other fractions chosen to represent the crude sample after the centrifugation (fraction 3), and fractions before (fraction 7) and after (fraction 21) those predicted to contain MVs in the gradient. We observed spherical, double membrane particles resembling MVs described in other Gram-positive species in fraction 15, but not in the other three fractions (**Figure 1B**) [2, 12]. To further characterize MVs observed in fraction 15, we combined fractions 13-16 in order to increase our sample mass, and measured the size and concentration of MVs by dynamic light scattering using a Nanosight NS300 instrument. MVs varied in size with a diameter ranging from 40-400 nm (**Figure S1**), similar to that reported for other Gram-positive bacteria [3, 13, 28]. To visualize MV formation in situ, we imaged live *E. faecalis* mounted on agar pads and stained with Nile Red in late-log phase. Acquired time series revealed increasing number of small (resolution limited) vesicles that appear to detach from the cell body, and continue to diffuse in close bacterial proximity, most likely trapped in a confined volume around the cell between agar and glass slide (**Figure 1C**, **Supplementary video 1A, 1B**).

### *E. faecalis* MVs are enriched in unsaturated lipids

The bacterial membrane is non-homogenous, containing distinct microdomains associated with functions including bacterial secretion and virulence, and which may serve as targets for antimicrobials [29, 30]. Since the *E. faecalis* membrane also contains lipid microdomains where secretion and virulence factor assembly functions are enriched [30–32] and since MVs are associated with distinct lipid repertoires in some bacteria [17], we considered the possibility that *E. faecalis* MVs are also released from specific lipid microdomains on the bacterial membrane. To address this question and ask whether *E. faecalis* MVs are composed of distinct lipid subsets, we analysed the lipidomes of purified MVs (25-30% of OptiPrep density) to entire membrane of late log phase bacterial cells. We analysed the lipid composition from five independent biological samples by liquid chromatography tandem mass spectrometry (LC-MS) using multiple reaction monitoring (MRM) methods previously established for (PG) and lysyl-phosphatidylglycerol (Lys-PG), as well as newly developed MRMs for diglucosyl-diacylglycerol (DGDAG) [24].

Within the PG species, which are the predominant lipid in the *E. faecalis* lipid membrane [24], we observed a significant increase in the overall abundance of polyunsaturated lipids (PG 32:2; PG 34:2), and reduction in monounsaturated and saturated lipids (PG 34:0, PG 35:0, PG 32:1, PG 34:1, PG 36:1) in MV samples as compared to whole cell lysates (WCL) (**Figure 2A**). For the less abundant DGDAG lipid species, the trend is similar in that we observe an increase in overall polyunsaturated species (**Figure 2B**). While we observed differences in individual species of Lys-PG (increased 34:1 and 33:2 in MVs, decreased 34:2 in MVs), the overall levels of saturation for Lys-PG were similar between MV and WCL. (**Figure 2C**).

**Figure 2.**
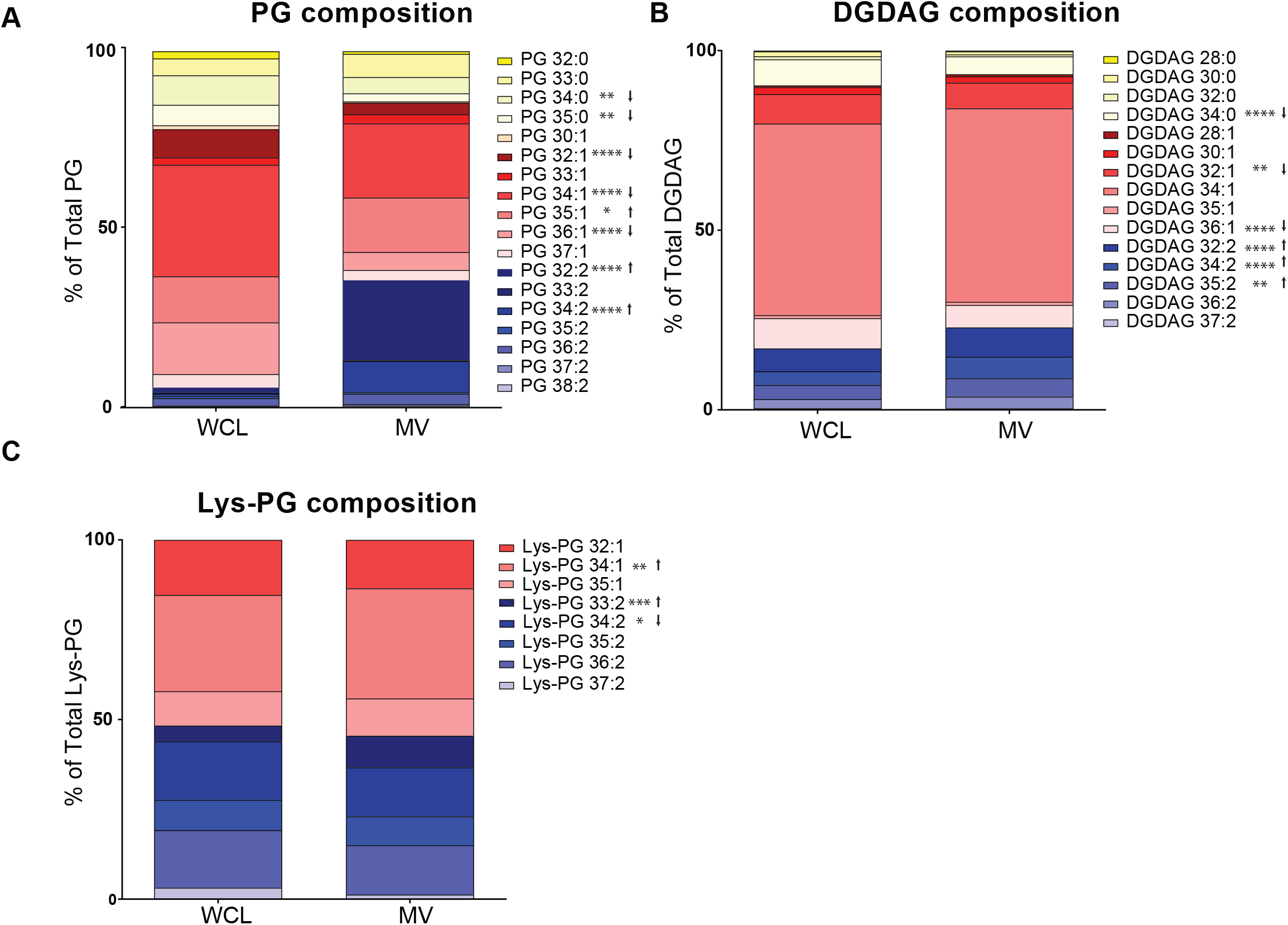
MVs are enriched in polyunsaturated PG species. Constituent distribution of individual lipid species from whole cell lysate (WCL) or MVs within the analysed lipid classes (**A**) phosphatidylglycerol (PG), (**B**) diglucosyl-diacylglycerol (DGDAG), and (**C**) lysyl-phosphatidylglycerol (Lys-PG). Each stack represents the mean from 6 biological replicates. *, P ≤ 0.05; **, P ≤ 0.01; ****, P ≤ 0.0001; Fisher’s LSD test for one-way ANOVA.

### *E. faecalis* MVs possess a unique proteome

To further characterize the properties of *E. faecalis* MVs, we performed proteomic analysis on OptiPrep-purified MVs and from late-log phase whole cells using LC/MS/MS [33]. Proteins for which three or more unique peptides were identified were considered in the analysis. In total, we identified 225 proteins from MV samples and 397 proteins from whole cell lysates, and sorted the proteins based on their calculated average abundance score (**Tables S4** and **S5**). Abundance was calculated as the number of unique peptides for a given protein divided by the total number of peptides within each sample, and the average abundance for each protein in the three biological replicates is reported. In addition, enrichment of each protein was calculated as the average abundance in the MV fraction divided by the average abundance of the same protein in the whole cell fraction. Next, we sorted the 15 most abundant and 15 enriched proteins in the MVs, and established 10 “signature” MV proteins that were both most abundant and most enriched in MV fractions relative to whole cells (**Figure 3, Table 1).** Within those are IreK (OG1RF_12384), a eukaryote-like Ser/Thr kinase and phosphatase; penicillin binding proteins Pbp1A, Pbp1B and PenA; a S1 family extracellular protease OG1RF_12235; lipoprotein (OG1RF_12508); CnaB and 2 ABC superfamily ATP binding cassette transporters (OG1RF 10124 and OG1RF_12508).

**Figure 3.**
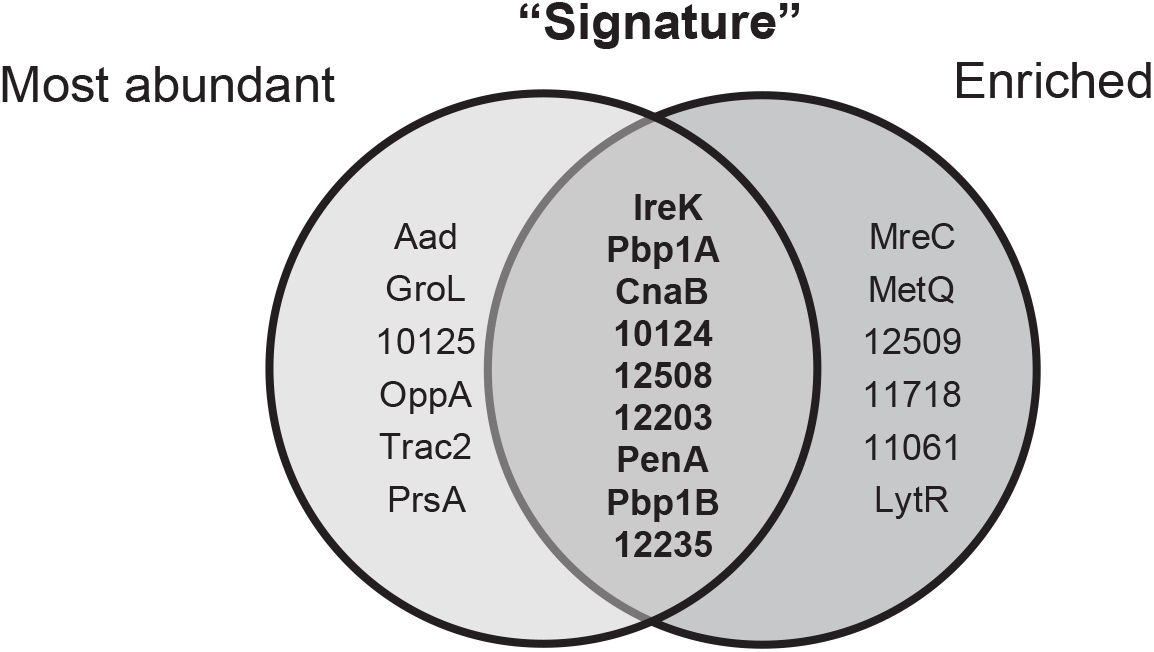
MVs possess a unique protein profile with enrichment and abundance of “signature” proteins. The Venn diagram shows the top 15 most abundant and enriched proteins in the MV fraction, by annotation where available or by gene number. Abundance was calculated as the number of unique peptides/total number of peptides. Enrichment was calculated as abundance in MVs/abundance in WCL. The gene description and position within each fraction are listed in **Table 1** in descending order based on the rank of abundance.

**Table 1.** Function and rank position of the top 15 most abundant and enriched proteins identified in MVs by MS. Where n/p – not present in WCL within 397 identified proteins; in bold are the most abundant and the most enriched “signature” proteins (description to **Figure 3**).

| Protein | Position in MVs (of 225) | Position in WCL (of 397) | Function |
| --- | --- | --- | --- |
| Aad | 1 | 8 | Aldehyde-alcohol dehydrogenase |
| GroL | 2 | 15 | Chaperonin |
| **OG1RF_12384** | **3** | **168** | **Non-specific serine/threonine protein kinase** |
| **Pbp1A** | **4** | **n/p** | **Penicillin binding proteins 1A** |
| **CnaB** | **5** | **n/p** | **Cna protein B-type domain protein** |
| **OG1RF_10124** | **6** | **151** | **ABC superfamily ATP binding cassette transporter, binding protein** |
| **OG1RF_12508** | **7** | **n/p** | **Thiamine biosynthesis lipoprotein** |
| **OG1RF_12203** | **8** | **155** | **ABC superfamily ATP binding cassette transporter, substrate-binding protein** |
| OG1RF_10125 | 9 | 67 | ABC superfamily ATP binding cassette transporter, binding protein |
| **PenA** | **10** | **n/p** | **Penicillin-binding protein 2 gene** |
| OppA | 11 | 33 | Oligopeptide ABC superfamily ATP binding cassette transporter, binding protein |
| TraC2 | 12 | 93 | Peptide ABC superfamily ATP binding cassette transporter, binding protein |
| **OG1RF_12235** | **13** | **n/p** | **S1 family extracellular protease** |
| **Pbp1B** | **14** | **n/p** | **Penicillin-binding protein 1B** |
| PrsA | 15 | 118 | Peptidyl-prolyl cis-trans isomerase |
| MreC | 19 | n/p | Rod shape-determining protein MreC |
| MetQ | 20 | n/p | ABC superfamily ATP binding cassette transporter, binding protein |
| OG1RF_12509 | 23 | 78 | Pheromone cAD1 lipoprotein |
| OG1RF_11718 | 24 | 220 | Hypothetical protein |
| OG1RF_11061 | 26 | n/p | Phage tail protein |
| LytR | 32 | n/p | Response regulator |

In addition to a phage tail protein OG1RF_11061 that was found among the 15 most enriched proteins, we detected an additional 4 phage proteins encoded at a previously described phage02 locus that are only present in MV fraction and are not found in a whole cell lysate (**Table S4**). The phage02-encoding operon (OG1RF 11046-11063) is a part of the *E. faecalis* core genome and considered cryptic because it lacks crucial genes for DNA packaging and excision [34]. However, we observed phage tail-like structures in the purified MV fraction by TEM (**Figure 4A**). In *Pseudomonas aeruginosa* MVs can be formed as a result of explosive cell lysis mediated by cell-wall modifying phage endolysins that degrade the bacterial membrane to burst the bacterial cell, resulting in MV formation and spontaneous membrane reannealing [7]. We hypothesized that phage tails could be assembled inside *E. faecalis* cells and released through explosive cell lysis, concurrently contributing to MV formation. We performed TEM on mid-log phase bacteria and observed MVs closely associated with the bacterial surface (**Figure 4B**), similar to bacteria-associated MVs observed in *P. aeruginosa* upon phage-mediated cell lysis [7]. To test whether phage-mediated explosive cell lysis occurred in *E. faecalis*, we first generated a strain in which the entire phage02 operon was deleted (Δ*pp2*), including the predicted phage structural proteins, holin, and endolysin genes. We performed TEM on MVs isolated from WT and Δ*pp2* and confirmed absence of the phage tails in the Δ*pp2* phage deletion mutant (data not shown).

**Figure 4.**
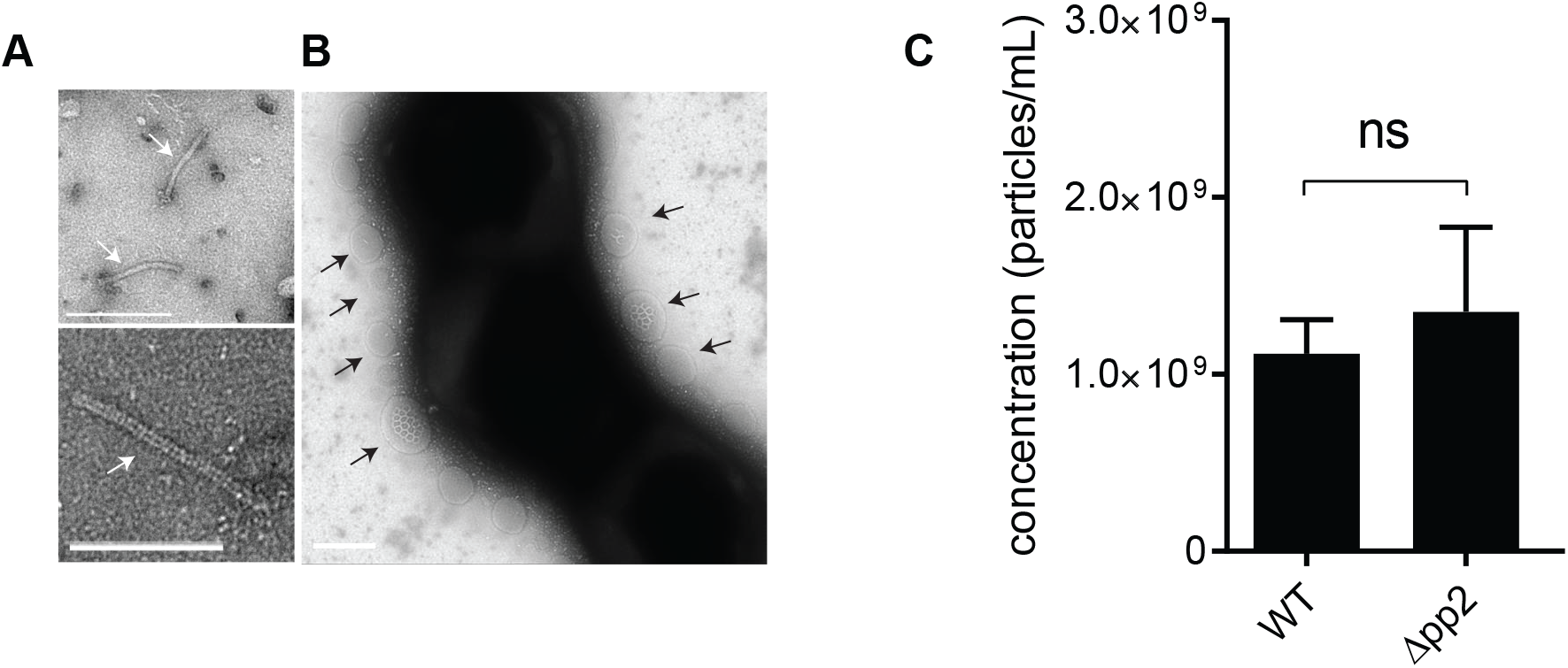
Phage tails co-purify with MVs, but phage tail production does not contribute to MV abundance. (**A**) TEM on assembled phage tails that are present within purified MVs. Scale bar is 100 nm. (**B**) TEM image of negatively stained *E. faecalis*, where MVs are associated with the cells surface. Scale bar is 100 nm. (**C**) The concentration of MVs in WT and the Δ*pp2* mutant as determined by Nanosight from 3 independent experiments. Statistical analysis was performed by the unpaired t-test. ns: P>0.05.

To determine whether the phage02-encoded factors, such as an endolysin, contributed to cell lysis and MV formation, we quantified MVs from WT and Δ*pp2*, but observed no difference in MV numbers between the two strains (**Figure 4C**). Since phage02 is the only predicted phage element in the OG1RF genome, these findings suggested that a phage-mediated mechanism, such explosive cell lysis, does not significantly contribute to *E. faecalis* MV formation.

### *E. faecalis* MVs do not mediate intra-species or cross-genera horizontal gene transfer *in vitro*

OMVs from Gram-negative bacteria may contain DNA from chromosomal, plasmid, or viral origin [35]. Plasmid and chromosomal DNA carrying antibiotic resistance genes can be transferred via OMVs to cells of the same or different genera to confer antibiotic resistance in the recipient [36, 37]. Among Gram-positive bacteria, *Clostridium perfringens* and *S. mutants* pack chromosomal DNA in MVs [11, 38]. A single report on MV-mediated horizontal gene transfer (HGT) in Gram-positive bacteria demonstrated the ability of a MV-containing fraction from WT *Ruminococcus sp.* strain YE71 to transform and permanently restore cellulose-degrading activity to a Cel^−^ mutant that was otherwise unable to degrade cellulose [39]. We tested if *E. faecalis* can pack plasmids inside MVs for HGT to another *E. faecalis* strain or to another species. We first isolated MVs from plasmid-carrying OG1RF pGCP123 [22], a 3045 bp, non-conjugative plasmid that can stably replicate in both Gram-positive and Gram-negative bacteria and which encodes kanamycin-resistance.

To first determine if *E. faecalis* MVs contain DNA, we quantified DNA from the intact MV prep before and after the MV lysis. We observed a ~150% increase in DNA concentration from lysed sample, compared to the intact MV, indicating that bacterial DNA is present within MV lumen (**Figure 5A**). To determine if plasmid DNA is present within the MV prep, we performed PCR with plasmid-specific primers on the crude and purified MV fractions, using purified pGCP123 plasmid as a control. We observed a plasmid-specific PCR product of 229 bp in the plasmid control and MV sample from plasmid-carrying strains (**Figure 5B**) indicating that plasmid DNA is present in both crude and purified MV fractions. In principle, plasmid DNA can be either packed inside MVs or associated with the exterior of MVs (and therefore co-purified with MVs). To address localization of pGCP123 with respect to *E. faecalis* MVs, we exposed MVs to DNase in order to degrade any extracellular plasmid (**Figure 5C**). To control for efficiency of DNase treatment, we used purified plasmid and lysed MVs as controls. We observed a plasmid-specific PCR product only in DNase untreated samples. To ensure that DNase was fully inactivated after incubation with MVs, and didn’t continue to digest plasmid DNA released from MVs during the PCR reaction, we mixed purified MVs with the same amount of DNase followed by immediate DNase inactivation by boiling at 98°C for 10 min. We observed a plasmid-specific PCR product in the inactivated DNase sample suggesting that DNase was fully inactivated before the PCR reaction. Hence, the observed an increase in DNA concentration upon MV lysis (**Figure 5A**), indicates that bacterial DNA is present within MVs. This MV resident DNA, however, likely does not contain plasmids since we show that plasmid DNA is not present within MV lumen, but instead co-purifies with MVs.

**Figure 5.**
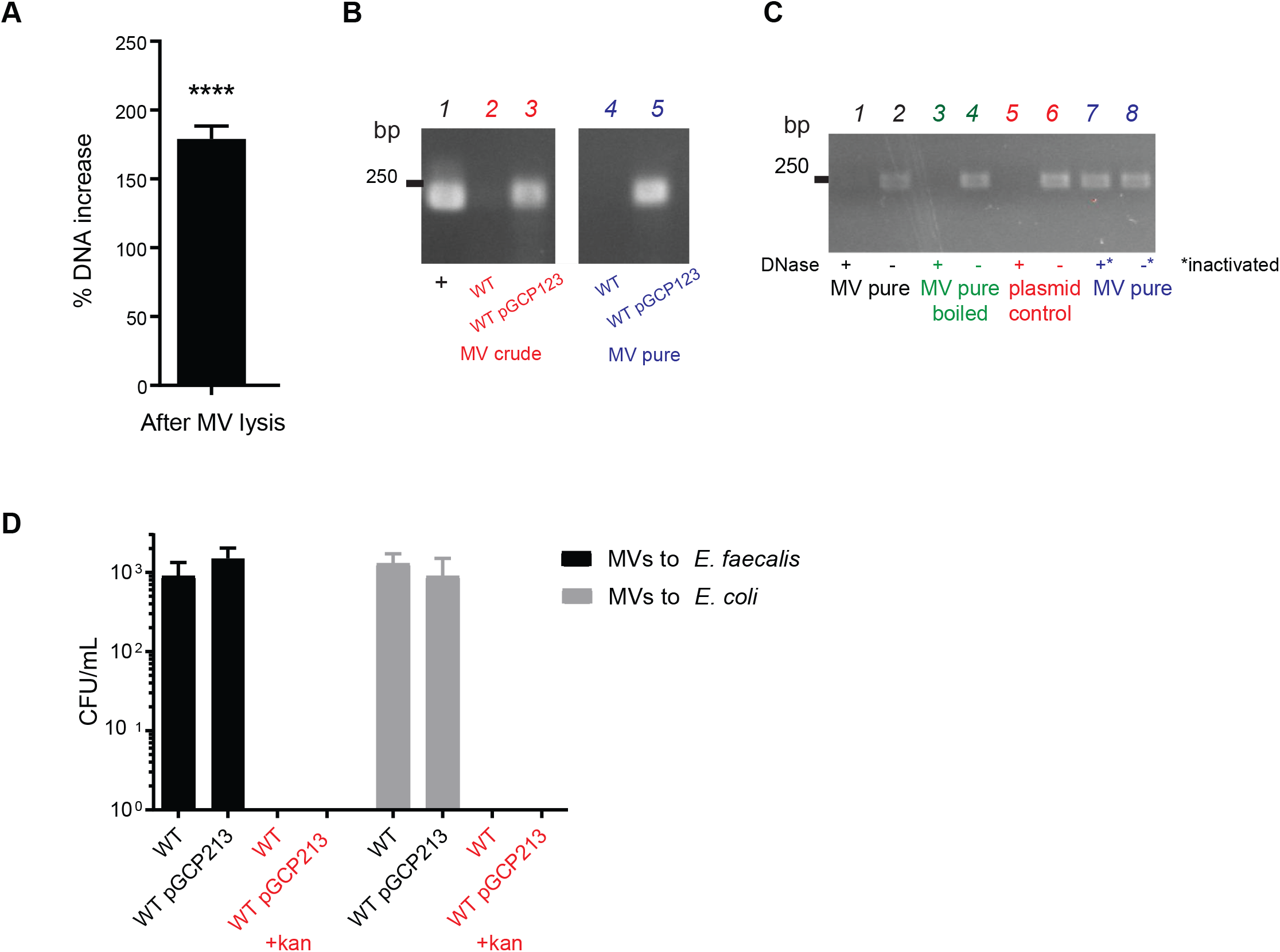
Plasmid DNA co-purifies with MVs but is not transferred to *E. faecalis* or *E. coli* cells. (**A**) DNA concentration was measured by Qubit in intact MV samples and in MVs lysed by boiling. Data shown from 3 independent experiments. Statistical analysis was performed by the unpaired t-test using GraphPad. ****, p<0.0001. (**B**) Agarose gel showing PCR product amplified with plasmid-specific primers on crude and purified MV fractions of WT and WT pGCP123. The expected plasmid PCR product is 240 bp. (**C**) Agarose gel showing PCR product amplified with plasmid-specific primers on intact and lysed MVs from WT pGCP123, subjected to DNAse treatment or treatment with inactivated DNAse prior to PCR. DNAse treated pGCP123 serves as a control. (**D**) The number of transformants following *E. faecalis* and *E. coli* incubation with MVs extracted from WT or WT pGCP123 determined by CFU enumeration on non-selective BHI or selective media with kanamycin. Statistical analysis was performed by the one-way ANOVA using one-way ANOVA test with Tukey’s multiple comparison test. ****, p<0.0001.

To determine if plasmids co-purified with MVs could mediate HGT, we exposed purified MVs to plasmid free strains of *E. faecalis* and *E. coli* for 2 hours to allow for plasmid transfer prior plating bacteria on selective plates containing kanamycin. However, we detected no resistant colonies on the selective plates suggesting that plasmid was not transferred to *E. coli* or *E. faecalis* by MVs under these experimental conditions (**Figure 5D**).

### MVs modulate the NF-κB response in macrophages

Proteomic analysis identified 5 lipoproteins (OG1RF_11390, OG1RF_11130, OG1RF_11506, OG1RF_12508, OG1RF_12509) enriched in MVs, similar to reports of lipoprotein-rich MVs from *S. pyogenes* and *M. tuberculosis* [12, 16] (**Tables S3** and **S4**). Mycobacteria-derived MVs bear 2 lipoproteins which are demonstrated agonists of the Toll-like 2 receptor and that activate an inflammatory response in mice. By contrast, MVs from *S. pyogenes* do not activate TLR2, indicating that the immunogenic potential of MVs varies between species [12, 16, 40]. We have previously shown that *E. faecalis* activates macrophages at low multiplicities of infection (MOI) and is immunosuppressive to macrophages at high MOI [41]. To determine whether *E. faecalis* MVs contribute to immune suppression or activation, we tested the NF-κB response of macrophages upon exposure to *E. faecalis* MVs. We purified MVs using OptiPrep and added them to RAW-blue macrophages (1000 MVs/macrophage) with or without LPS to assess their ability to activate NF-κB signalling on their own, or suppress LPS-mediated activation. Purified lipopolysaccharide (LPS) served as a positive control for maximal NF-κB activation and 25% OptiPrep media as a negative control. We also combined the first 10 fractions from the OptiPrep gradient after centrifugation to serve as a MV-free concentrated supernatant for a secondary negative control. In this assay, we observed that *E. faecalis* MVs activate NF-κB signalling in macrophages (**Figure 6**). Since we observed that the MV fraction from WT *E. faecalis* co-purified with bacteriophage tails (**Figure 4**), and since fully assembled bacteriophages can be immunomodulatory and suppress phagocytosis and LPS-induced phosphorylation of NF-κBp65 [42–44], we next dissected whether phage tails contribute to MV-mediated activation of NF-κB in macrophages (**Figure 6**). We isolated MVs from Δ*pp2* lacking the whole prophage operon and compared the NF-κB response to the response elicited by MVs from WT. We observed no statistical difference in NF-κB reporter activity of Δ*pp2* mutant (MVs only) compared to WT (MVs + phage tails), suggesting that MVs alone are immunostimulatory and can induce NF-κB signalling in macrophages, regardless of the presence of co-purified phage tails (**Figure 6**).

**Figure 6.**
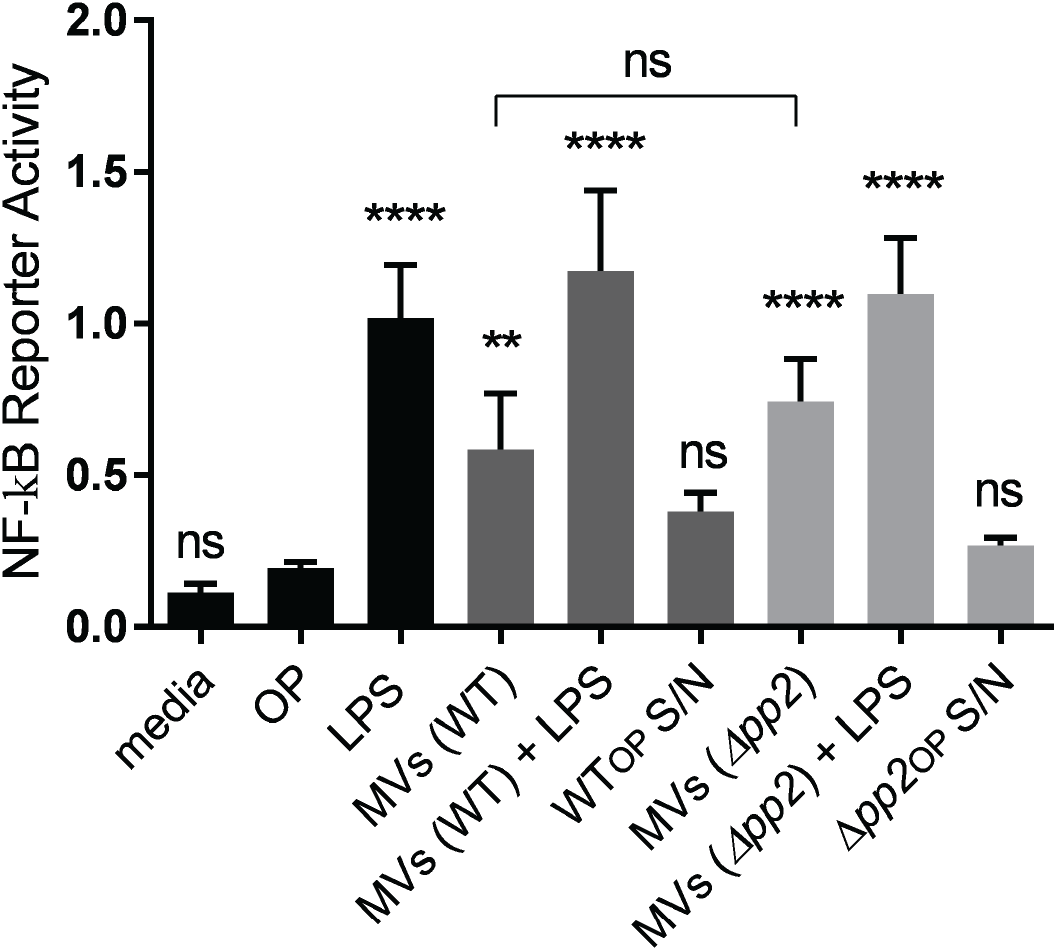
MVs but not phage tails activate NF-kB pathway in macrophages. RAW-blue cells derived from RAW267.4 macrophages were stimulated with lipopolysaccharide (LPS) at 100 ng/mL (positive control), OptiPrep (OP) (negative control), MV-free concentrated supernatant in OptiPrep from WT and Δ*pp2* (WT_OP_ S/N and Δ*pp2*_*OP*_ *S/N)* (secondary controls), MVs derived from WT and Δ*pp2* at 1000 particles/macrophage, and LPS + MVs. Six hours after stimulation, the NF-kB response was measured by secreted embryonic alkaline phosphatase reporter activity, transcribed from the plasmid under NF-kB inducible promoter. Statistical analysis was performed by the one-way ANOVA using one-way ANOVA test with Tukey’s multiple comparison test. ****, p<0.0001, ** p<0.001, ns: P>0.05 among all of the conditions as compared to OptiPrep negative control.

## Discussion

Vesicle shedding from bacteria is a common process in Gram-negative and Gram-positive bacteria. Yet the mechanisms of MV formation and function, especially in Gram-positive species are largely unknown. MVs of Gram-positive bacteria possess unique proteomes, are immunogenic, and in some cases contribute to host cells death by carrying toxins [2, 3, 40, 45]. In this study, we report that the opportunistic pathogen *E. faecalis* strain OG1RF produces MVs ranging in size from 40-400nm. *E. faecalis* MVs are enriched for a unique subset of proteins, unsaturated lipids, and are capable of activating NF-kB signalling in macrophages.

Several studies have compared the lipid profile of MVs to that of the complete bacterial membrane. In Group A streptococcus, MVs are enriched in PG and significantly reduced in CL while the saturation levels of fatty acids remained unchanged [17]. In *Propionibacterium acnes*, MVs also possess a distinct lipid profile with levels of triacylglycerol significantly lower compared to the cell membrane [46]. Both reports suggest that lipid composition and distribution in MVs can be very different from that of the bacterial cell membrane, supporting the hypothesis that MV formation is not a random process. In *E. faecalis*, we observed a significant increase in the levels of unsaturated PG in MVs as compared to the whole cell membrane. Unsaturated lipids enhance membrane fluidity and partition to ordered domains in more fluid regions of the bilayer [46, 47]. In model membranes, more flexible unsaturated lipids become concentrated in tubular regions pulled from a vesicle and unsaturated lipids are sorted into pathways involving highly curved tubular intermediates [48]. Therefore, *E. faecalis* microdomains with polyunsaturated PGs might provide additional flexibility for vesicle formation.

In addition to a distinct lipid profile, *E. faecalis* MVs possess a unique proteome. Among MV signature proteins are penicillin binding proteins, Pbp1A and Pbp1B, which dynamically localize in the inner membrane in *E. coli* but are also enriched in the septal divisome during cell division [49, 50]. In *Streptococcus pneumoniae* division and are localized to the septum region [51] These observations are consistent with a model in which *E. faecalis* MVs might be formed from the Pbp-enriched septal region during cell division. Consistent with this hypothesis, within the signature MV proteins, we detected the serine-threonine kinase IreK that monitors cell wall integrity and mediates adaptive responses to cell wall-active antibiotics [52, 53]. An IreK homologue in *S. pneumoniae* – StkP - is localized at the division septum via penicillin-binding protein and serine/threonine kinase associated (PASTA) domains linked to un-crosslinked PG, the same unique domains that are present in enterococcal IreK [54, 55]. The abundance of likely septum-localized proteins within *E. faecalis* MVs suggests that MV formation is a spatiotemporally organized process, and might be driven through IreK signalling at the septum. Moreover, by *in situ* live imaging MVs appeared to be derived from the septal region, further supporting septal vesiculation model.

Supplementary to MV release from septal polyunsaturated PG microdomains, it was formally possible that *E. faecalis* MVs might be formed through explosive cell lysis mediated by phage tail release, similar to MV formation during bacteriophage release in *P. aeruginosa* [7]. Indeed, we observed that MVs co-purified with fully-assembled phage tails encoded by a cryptic phage. While we didn’t observe significantly changed MV numbers between the WT and Δ*pp2* phage mutant, the question of what function these phage tails might confer to *E. faecalis* now stands. Moreover, in *P. aeruginosa*, a population of MVs were observed to be attached to cells adjacent to those which underwent lysis, which would not be reflected in MV quantification experiments. Therefore, we cannot yet rule out partial contribution of explosive cell lysis to MV formation in *E. faecalis*.

Finally, we observed that lipoproteins are enriched in MVs, similar to MVs from *S. pyogenes* and *M. tuberculosis* [12, 16]. Lipoproteins are potent inducers of the host inflammatory responses through TLR2 receptors, and lipoprotein-rich MVs from *M. tuberculosis* and *C. perfringens* activate macrophages, leading to the release of inflammatory cytokines [28, 40, 56]. In *L. monocytogenes,* pheromone cAD1, the homologue of pheromone cAD1 lipoprotein that is enriched in enterococcal MVs, enhances bacterial escape from host cell vacuoles and bacterial virulence [57]. We hypothesize that *E. faecalis* lipoprotein-enriched MVs may contribute to their ability to promote NF-kB activation in macrophages. In addition, it is possible that MV-associated unmethylated prokaryotic DNA is recognized by TLR9 receptors on host immune system cells leading to NF-kB activation [58, 59]. Finally, bacterial lipids can engage pattern recognition receptors on host cell membranes to control inflammation and immunity by interacting with TLR2 and TLR4 receptors [60, 61]. Whether lipids present within enterococcal MVs play a role in immune modulation remains unclear. The precise roles for enterococcal MVs during infection, whether they confer benefits via immune modulation or promoting bacterial growth and survival, are the subject of ongoing investigation.

In summary, we identified that *E. faecalis* MVs are composed of distinct protein and lipid profiles, suggesting they may arise via a regulated MV biogenesis process from the septum, where the cell-wall is thinnest and which may be enriched with more flexible polyunsaturated microdomains. Functionally, *E. faecalis* MVs activate NF-kB signalling in macrophages, possibly due to their abundance of immunogenic lipoproteins. Future work on mechanisms of MV formation and comparing the function of MVs from both commensal and pathogenic *E. faecalis* strains will enhance our understanding in enterococcal pathogenesis and host-pathogen interactions.

## Acknowledgements

This work was supported by the National Research Foundation and Ministry of Education Singapore under its Research Centre of Excellence Programme, as well as the National Research Foundation under its Singapore NRF Fellowship programme (NRF-NRFF2011-11). This work was also supported by a Tier 1 grant sponsored by the Singapore Ministry of Education (MOE2017-T1-001-269).

We are grateful to Jenny Dale and Gary Dunny for supplying us with *E. faecalis* OG1RF transposon mutants used in this study. We also thankful to the Taplin Biological Mass Spectrometry Facility and Ross Tomaino for the MS analysis. The TEM work was undertaken at the NTU Institute of Structural Biology Cryo-EM lab in Nanyang Technological University, Singapore. We thank Andrew Wong for the assistance in sample prep and imaging. We also thank Annika Sjöström and Monica Persson from Umea University, Sweden for advice on MV isolation and atomic force microscopy.

