## Supplementary Figure 1 for "The composition and function of *Enterococcus faecalis* membrane vesicles"

**Figure S1***.* **Size distribution and concentration of MVs in combined OptiPrep fractions 13-16 by Nanosight.** *E. faecalis* produces MVs ranging from 40 to 400nm in size. Distinct peaks represent a specific particle size range. Data shown are combined from 3 independent experiments, where the back line is the mean and the grey shaded area is a standard deviation. Particle concentration was calculated automatically by Nanosight software.

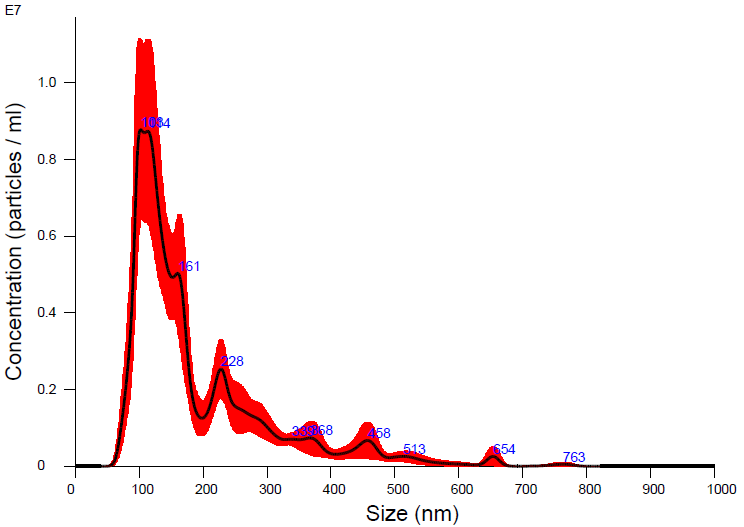
