## Supplementary Table 1 for "The composition and function of *Enterococcus faecalis* membrane vesicles"

**Table S1. Primers used in the study.**

| **Primer name** | **Sequence (5`-3`)** |
| --- | --- |
| pp2.infu.1F | TGTAAAACGACGGCCAGTGAGTTCGCCCAACACCACAAGC |
| pp2.infu.1R | GCAAAATCTCTGCTGAATGCCCGACGCCAAGCATAAG  ATTTGG |
| pp2.infu.2F | AATCTTATGCTTGGCGTCGGGCATTCAGCAGAGATTTT  GCCA |
| pp2.infu.2R | CAGGAAACAGCTATGACCATCATGCGTTGGGTGTGGCA |
| pgcp213.infu.F | ATGGTCATAGCTGTTTCCTGTGTG |
| pgcp213.infu.R | TCACTGGCCGTCGTTTTACAAC |
| infu_check_pp2F | GGTTTTGAATGTGGCGCTAT |
| infu_check_pp2R | AGTGTCCGCACATAGGTTCC |
