## Supplementary Table 2 for "The composition and function of *Enterococcus faecalis* membrane vesicles"

**Table S2. Standards used for lipidomic analysis.** Lipid standards obtained from Avanti Polar Lipids, USA.

| Standard | Usage | Catalogue # | Stock conc. [mg/mL] | Final conc. [µg/mL] | Precursor ion (m1) m/z | Fragment ion (m3) m/z |
| --- | --- | --- | --- | --- | --- | --- |
| PG 14:0 | Internal | 840445P | 1 | 5 | 665.5 [M-H]^-^ | 153 |
| Lys-PG 16:0 | Internal | 840520P | 0.1 | 4 | 849.6 [M-H]^-^ | 145 |
| MGDAG 34:1 | External | 840522P | 1 | variable | 774.6 [M+NH_4_]^+^ | 313 |
