## Supplementary Table 3 for "The composition and function of *Enterococcus faecalis* membrane vesicles"

**Table S3. MRM transitions list for DGDAG and MGDAG**

| Lipid Species | Ion Mode | Precursor Ion | Product Ion |
| --- | --- | --- | --- |
| MGDAG 34:1 | Positive | 774.6 | 313.3 |
| DGDAG 28:1 | Positive | 852.6 | 493.5 |
| DGDAG 28:0 | Positive | 854.6 | 495.5 |
| DGDAG 30:1 | Positive | 880.6 | 521.5 |
| DGDAG 30:0 | Positive | 882.6 | 523.5 |
| DGDAG 32:2 | Positive | 906.6 | 547.5 |
| DGDAG 32:1 | Positive | 908.6 | 549.5 |
| DGDAG 32:0 | Positive | 910.6 | 551.5 |
| DGDAG 34:2 | Positive | 934.6 | 575.5 |
| DGDAG 34:1 | Positive | 936.7 | 577.6 |
| DGDAG 34:0 | Positive | 938.7 | 579.6 |
| DGDAG 35:2 | Positive | 948.7 | 589.6 |
| DGDAG 35:1 | Positive | 950.7 | 591.6 |
| DGDAG 36:2 | Positive | 962.7 | 603.6 |
| DGDAG 36:1 | Positive | 964.7 | 605.6 |
| DGDAG 37:2 | Positive | 976.7 | 617.6 |
