## Supplementary Table 4 for "The composition and function of *Enterococcus faecalis* membrane vesicles"

**Table S4**. **Proteins identified in whole cell lysate by LC/MS/MS.** Proteins listed in descending order based on the abundance score. Abundance (A*) score is calculated as a number of unique peptides for each protein vs the number of total peptides within the sample.

| A* position | protein name | A*1 | A*2 | A*3 | A*_avr_ |
| --- | --- | --- | --- | --- | --- |
| 1 | rpoC | **1.423** | **1.398** | **1.439** | **1.420** |
| 2 | fusA | 1.220 | 1.158 | 1.474 | 1.284 |
| 3 | tuf | 0.929 | 1.114 | 1.790 | 1.278 |
| 4 | dnaK | 0.900 | 1.049 | 1.334 | 1.094 |
| 5 | rpoB | 1.016 | 1.092 | 1.053 | 1.054 |
| 6 | eno | 0.842 | 0.874 | 1.299 | 1.005 |
| 7 | pflB | 0.755 | 1.027 | 1.123 | 0.968 |
| 8 | aad | 0.842 | 0.961 | 0.983 | 0.929 |
| 9 | clpB | 0.871 | 0.852 | 1.018 | 0.914 |
| 10 | pgk | 0.900 | 0.786 | 0.913 | 0.866 |
| 11 | tig | 0.813 | 0.874 | 0.842 | 0.843 |
| 12 | ptsI | 0.784 | 0.808 | 0.878 | 0.823 |
| 13 | infB | 0.871 | 0.721 | 0.807 | 0.800 |
| 14 | pyk | 0.697 | 0.743 | 0.948 | 0.796 |
| 15 | groL | 0.726 | 0.699 | 0.948 | 0.791 |
| 16 | tsf | 0.639 | 0.765 | 0.913 | 0.772 |
| 17 | OG1RF_11938 | 0.784 | 0.590 | 0.913 | 0.762 |
| 18 | guaA | 0.697 | 0.633 | 0.807 | 0.713 |
| 19 | secA | 0.552 | 0.786 | 0.702 | 0.680 |
| 20 | clpE | 0.581 | 0.633 | 0.737 | 0.650 |
| 21 | ftsZ | 0.610 | 0.633 | 0.702 | 0.648 |
| 22 | tyrS | 0.552 | 0.590 | 0.772 | 0.638 |
| 23 | pgi | 0.494 | 0.590 | 0.772 | 0.619 |
| 24 | tkt | 0.639 | 0.612 | 0.597 | 0.616 |
| 25 | pnp | 0.552 | 0.590 | 0.702 | 0.614 |
| 26 | glnP | 0.552 | 0.590 | 0.667 | 0.603 |
| 27 | rpoA | 0.552 | 0.568 | 0.667 | 0.596 |
| 28 | glmS | 0.581 | 0.481 | 0.702 | 0.588 |
| 29 | typA | 0.610 | 0.546 | 0.597 | 0.584 |
| 30 | ftsH | 0.610 | 0.633 | 0.491 | 0.578 |
| 31 | rplE | 0.465 | 0.524 | 0.737 | 0.575 |
| 32 | OG1RF_11602 | 0.697 | 0.677 | 0.351 | 0.575 |
| 33 | oppA | 0.581 | 0.546 | 0.562 | 0.563 |
| 34 | lysS | 0.639 | 0.568 | 0.456 | 0.554 |
| 35 | alaS | 0.668 | 0.502 | 0.491 | 0.554 |
| 36 | OG1RF_12374 | 0.581 | 0.568 | 0.491 | 0.547 |
| 37 | trxB2 | 0.465 | 0.590 | 0.562 | 0.539 |
| 38 | ileS | 0.581 | 0.502 | 0.527 | 0.537 |
| 39 | srmB | 0.523 | 0.546 | 0.527 | 0.532 |
| 40 | gap2 | 0.407 | 0.459 | 0.702 | 0.522 |
| 41 | argS | 0.523 | 0.655 | 0.351 | 0.510 |
| 42 | metG | 0.523 | 0.546 | 0.456 | 0.508 |
| 43 | ndh3 | 0.523 | 0.437 | 0.562 | 0.507 |
| 44 | rplB | 0.407 | 0.481 | 0.632 | 0.506 |
| 45 | gnd | 0.523 | 0.415 | 0.562 | 0.500 |
| 46 | rpsD | 0.407 | 0.437 | 0.632 | 0.492 |
| 47 | OG1RF_10367 | 0.581 | 0.459 | 0.421 | 0.487 |
| 48 | pgcA | 0.465 | 0.481 | 0.456 | 0.467 |
| 49 | rpsA | 0.377 | 0.437 | 0.562 | 0.459 |
| 50 | ackA | 0.552 | 0.437 | 0.386 | 0.458 |
| 51 | ccpA | 0.494 | 0.371 | 0.491 | 0.452 |
| 52 | ndh | 0.465 | 0.393 | 0.491 | 0.450 |
| 53 | atpD2 | 0.465 | 0.459 | 0.386 | 0.436 |
| 54 | aceF | 0.407 | 0.371 | 0.527 | 0.435 |
| 55 | ldh | 0.436 | 0.306 | 0.562 | 0.434 |
| 56 | guaB | 0.407 | 0.437 | 0.456 | 0.433 |
| 57 | OG1RF_12366 | 0.494 | 0.415 | 0.386 | 0.432 |
| 58 | glyS | 0.523 | 0.481 | 0.281 | 0.428 |
| 59 | ezrA | 0.494 | 0.437 | 0.351 | 0.427 |
| 60 | divIVA | 0.436 | 0.350 | 0.491 | 0.425 |
| 61 | valS | 0.348 | 0.546 | 0.351 | 0.415 |
| 62 | fbaB | 0.348 | 0.415 | 0.456 | 0.407 |
| 63 | proS | 0.494 | 0.437 | 0.281 | 0.404 |
| 64 | manX2 | 0.348 | 0.371 | 0.491 | 0.404 |
| 65 | pdhA | 0.348 | 0.371 | 0.491 | 0.404 |
| 66 | OG1RF_10125 | 0.465 | 0.393 | 0.351 | 0.403 |
| 67 | rpsE | 0.377 | 0.328 | 0.491 | 0.399 |
| 68 | npr | 0.377 | 0.350 | 0.456 | 0.394 |
| 69 | rplA | 0.377 | 0.350 | 0.456 | 0.394 |
| 70 | gatB | 0.348 | 0.502 | 0.316 | 0.389 |
| 71 | glnA | 0.407 | 0.371 | 0.386 | 0.388 |
| 72 | glyA | 0.523 | 0.393 | 0.246 | 0.387 |
| 73 | pfkA | 0.377 | 0.328 | 0.456 | 0.387 |
| 74 | rpsG | 0.290 | 0.328 | 0.527 | 0.382 |
| 75 | pdhB | 0.465 | 0.328 | 0.351 | 0.381 |
| 76 | atpA2 | 0.348 | 0.371 | 0.421 | 0.380 |
| 77 | OG1RF_12509 | 0.348 | 0.371 | 0.421 | 0.380 |
| 78 | pta | 0.436 | 0.284 | 0.421 | 0.380 |
| 79 | nrdE | 0.465 | 0.459 | 0.211 | 0.378 |
| 80 | gatA | 0.377 | 0.437 | 0.316 | 0.377 |
| 81 | rpoD | 0.348 | 0.393 | 0.386 | 0.376 |
| 82 | ligA | 0.348 | 0.262 | 0.491 | 0.367 |
| 83 | upp2 | 0.377 | 0.371 | 0.351 | 0.367 |
| 84 | pheT | 0.319 | 0.393 | 0.386 | 0.366 |
| 85 | arcB | 0.436 | 0.306 | 0.351 | 0.364 |
| 86 | rplC | 0.319 | 0.350 | 0.421 | 0.363 |
| 87 | lpd | 0.523 | 0.459 | 0.105 | 0.362 |
| 88 | gltX | 0.319 | 0.371 | 0.386 | 0.359 |
| 89 | pepS | 0.407 | 0.350 | 0.316 | 0.357 |
| 90 | pyrG | 0.348 | 0.350 | 0.351 | 0.350 |
| 91 | rplF | 0.319 | 0.306 | 0.421 | 0.349 |
| 92 | traC2 | 0.377 | 0.350 | 0.316 | 0.348 |
| 93 | rpsB | 0.348 | 0.328 | 0.351 | 0.342 |
| 94 | opuAA | 0.407 | 0.371 | 0.246 | 0.341 |
| 95 | rplJ | 0.232 | 0.284 | 0.491 | 0.336 |
| 96 | fabF | 0.261 | 0.350 | 0.386 | 0.332 |
| 97 | dnaN | 0.319 | 0.350 | 0.316 | 0.328 |
| 98 | pyrH | 0.290 | 0.306 | 0.386 | 0.327 |
| 99 | OG1RF_11455 | 0.348 | 0.328 | 0.281 | 0.319 |
| 100 | ndh2 | 0.290 | 0.350 | 0.316 | 0.319 |
| 101 | fruA | 0.377 | 0.437 | 0.140 | 0.318 |
| 102 | leuS | 0.290 | 0.415 | 0.246 | 0.317 |
| 103 | ychF | 0.377 | 0.328 | 0.246 | 0.317 |
| 104 | OG1RF_11725 | 0.348 | 0.459 | 0.140 | 0.316 |
| 105 | OG1RF_10957 | 0.203 | 0.393 | 0.351 | 0.316 |
| 106 | rplW | 0.319 | 0.240 | 0.386 | 0.315 |
| 107 | fabI | 0.290 | 0.262 | 0.386 | 0.313 |
| 108 | rpsC | 0.232 | 0.284 | 0.421 | 0.312 |
| 109 | adk | 0.261 | 0.284 | 0.386 | 0.310 |
| 110 | clpX | 0.261 | 0.284 | 0.386 | 0.310 |
| 111 | engA | 0.319 | 0.328 | 0.281 | 0.309 |
| 112 | gyrA | 0.261 | 0.350 | 0.316 | 0.309 |
| 113 | ahpC | 0.348 | 0.262 | 0.316 | 0.309 |
| 114 | OG1RF_11970 | 0.319 | 0.284 | 0.316 | 0.306 |
| 115 | pepV | 0.261 | 0.262 | 0.386 | 0.303 |
| 116 | metK | 0.203 | 0.350 | 0.351 | 0.301 |
| 117 | prsA | 0.290 | 0.262 | 0.351 | 0.301 |
| 118 | glmM | 0.290 | 0.328 | 0.281 | 0.300 |
| 119 | fabD | 0.319 | 0.328 | 0.246 | 0.298 |
| 120 | accC | 0.319 | 0.218 | 0.351 | 0.296 |
| 121 | OG1RF_10418 | 0.319 | 0.284 | 0.281 | 0.295 |
| 122 | asnS | 0.203 | 0.328 | 0.351 | 0.294 |
| 123 | frr | 0.232 | 0.262 | 0.386 | 0.294 |
| 124 | clpP | 0.319 | 0.240 | 0.316 | 0.292 |
| 125 | OG1RF_11094 | 0.348 | 0.350 | 0.176 | 0.291 |
| 126 | rpsM | 0.232 | 0.284 | 0.351 | 0.289 |
| 127 | pepT | 0.290 | 0.328 | 0.246 | 0.288 |
| 128 | fabF2 | 0.290 | 0.218 | 0.351 | 0.287 |
| 129 | prfC | 0.319 | 0.284 | 0.246 | 0.283 |
| 130 | OG1RF_11278 | 0.261 | 0.262 | 0.316 | 0.280 |
| 131 | rpsH | 0.261 | 0.262 | 0.316 | 0.280 |
| 132 | OG1RF_11240 | 0.348 | 0.350 | 0.140 | 0.279 |
| 133 | hslU | 0.290 | 0.262 | 0.281 | 0.278 |
| 134 | fruK2 | 0.145 | 0.197 | 0.491 | 0.278 |
| 135 | rplO | 0.232 | 0.240 | 0.351 | 0.275 |
| 136 | prs1 | 0.261 | 0.240 | 0.316 | 0.273 |
| 137 | gdhA | 0.319 | 0.284 | 0.211 | 0.271 |
| 138 | OG1RF_10016 | 0.319 | 0.284 | 0.211 | 0.271 |
| 139 | purA | 0.319 | 0.284 | 0.211 | 0.271 |
| 140 | prfA | 0.261 | 0.306 | 0.246 | 0.271 |
| 141 | ftsA | 0.436 | 0.371 | 0.000 | 0.269 |
| 142 | pepF | 0.232 | 0.328 | 0.246 | 0.269 |
| 143 | rplQ | 0.261 | 0.218 | 0.316 | 0.265 |
| 144 | OG1RF_10963 | 0.319 | 0.262 | 0.211 | 0.264 |
| 145 | pepA2 | 0.290 | 0.218 | 0.281 | 0.263 |
| 146 | yhgF | 0.261 | 0.350 | 0.176 | 0.262 |
| 147 | rpsJ | 0.174 | 0.218 | 0.386 | 0.260 |
| 148 | lepA | 0.290 | 0.306 | 0.176 | 0.257 |
| 149 | ffh | 0.203 | 0.350 | 0.211 | 0.254 |
| 150 | OG1RF_10124 | 0.261 | 0.218 | 0.281 | 0.254 |
| 151 | pycA | 0.290 | 0.328 | 0.140 | 0.253 |
| 152 | folD | 0.203 | 0.197 | 0.351 | 0.250 |
| 153 | rplM | 0.145 | 0.218 | 0.386 | 0.250 |
| 154 | OG1RF_12203 | 0.290 | 0.350 | 0.105 | 0.248 |
| 155 | uvrA | 0.261 | 0.306 | 0.176 | 0.248 |
| 156 | OG1RF_11165 | 0.232 | 0.262 | 0.246 | 0.247 |
| 157 | rplK | 0.261 | 0.197 | 0.281 | 0.246 |
| 158 | rplS | 0.203 | 0.218 | 0.316 | 0.246 |
| 159 | OG1RF_10392 | 0.319 | 0.240 | 0.176 | 0.245 |
| 160 | thrS | 0.261 | 0.328 | 0.140 | 0.243 |
| 161 | greA | 0.232 | 0.175 | 0.316 | 0.241 |
| 162 | cap4C | 0.145 | 0.328 | 0.246 | 0.240 |
| 163 | fabG3 | 0.261 | 0.240 | 0.211 | 0.237 |
| 164 | OG1RF_12067 | 0.261 | 0.240 | 0.211 | 0.237 |
| 165 | nrdF | 0.232 | 0.197 | 0.281 | 0.237 |
| 166 | OG1RF_11464 | 0.261 | 0.306 | 0.140 | 0.236 |
| 167 | OG1RF_12384 | 0.261 | 0.306 | 0.140 | 0.236 |
| 168 | xpt | 0.319 | 0.175 | 0.211 | 0.235 |
| 169 | nrdD | 0.174 | 0.175 | 0.351 | 0.233 |
| 170 | nusA | 0.348 | 0.350 | 0.000 | 0.233 |
| 171 | mutS | 0.290 | 0.262 | 0.140 | 0.231 |
| 172 | zwf | 0.290 | 0.262 | 0.140 | 0.231 |
| 173 | glmU | 0.203 | 0.240 | 0.246 | 0.230 |
| 174 | OG1RF_12224 | 0.290 | 0.218 | 0.176 | 0.228 |
| 175 | atpA | 0.232 | 0.240 | 0.211 | 0.228 |
| 176 | sodA | 0.174 | 0.153 | 0.351 | 0.226 |
| 177 | arcA | 0.261 | 0.240 | 0.176 | 0.226 |
| 178 | pcrA | 0.232 | 0.218 |  | 0.225 |
| 179 | pepQ | 0.261 | 0.197 | 0.211 | 0.223 |
| 180 | rpmE | 0.261 | 0.197 | 0.211 | 0.223 |
| 181 | hup | 0.232 | 0.153 | 0.281 | 0.222 |
| 182 | clpC | 0.290 | 0.371 | 0.000 | 0.221 |
| 183 | dnaJ | 0.203 | 0.240 | 0.211 | 0.218 |
| 184 | oppF | 0.174 | 0.262 | 0.211 | 0.216 |
| 185 | rplT | 0.203 | 0.197 | 0.246 | 0.215 |
| 186 | tyrS | 0.203 | 0.197 | 0.246 | 0.215 |
| 187 | recA | 0.203 | 0.262 | 0.176 | 0.214 |
| 188 | cspA3 | 0.203 | 0.153 | 0.281 | 0.212 |
| 189 | OG1RF_10783 | 0.203 | 0.218 | 0.211 | 0.211 |
| 190 | rplN | 0.145 | 0.240 | 0.246 | 0.210 |
| 191 | rmlB | 0.232 | 0.218 | 0.176 | 0.209 |
| 192 | gyrB | 0.232 | 0.393 | 0.000 | 0.208 |
| 193 | aroF | 0.203 | 0.175 | 0.246 | 0.208 |
| 194 | murD | 0.203 | 0.240 | 0.176 | 0.206 |
| 195 | gpmA | 0.145 | 0.218 | 0.246 | 0.203 |
| 196 | serS | 0.232 | 0.197 | 0.176 | 0.201 |
| 197 | OG1RF_10977 | 0.319 | 0.284 | 0.000 | 0.201 |
| 198 | taq | 0.174 | 0.218 | 0.211 | 0.201 |
| 199 | rplD | 0.203 | 0.153 | 0.246 | 0.201 |
| 200 | rplI | 0.232 | 0.262 | 0.105 | 0.200 |
| 201 | polA | 0.290 | 0.306 | 0.000 | 0.199 |
| 202 | dps | 0.174 | 0.175 | 0.246 | 0.198 |
| 203 | purB | 0.203 | 0.175 | 0.211 | 0.196 |
| 204 | OG1RF_10072 | 0.232 | 0.175 | 0.176 | 0.194 |
| 205 | lplA2 | 0.290 | 0.284 | 0.000 | 0.191 |
| 206 | OG1RF_10021 | 0.174 | 0.153 | 0.246 | 0.191 |
| 207 | rpsK | 0.174 | 0.153 | 0.246 | 0.191 |
| 208 | thrC | 0.232 | 0.197 | 0.140 | 0.190 |
| 209 | phoU | 0.232 | 0.153 | 0.176 | 0.187 |
| 210 | efp | 0.203 | 0.175 | 0.176 | 0.185 |
| 211 | accD | 0.145 | 0.197 | 0.211 | 0.184 |
| 212 | mnmG | 0.203 | 0.240 | 0.105 | 0.183 |
| 213 | ppaC | 0.232 | 0.175 | 0.140 | 0.182 |
| 214 | hisS | 0.174 | 0.197 | 0.176 | 0.182 |
| 215 | OG1RF_10495 | 0.174 | 0.197 | 0.176 | 0.182 |
| 216 | rplL2 | 0.145 | 0.153 | 0.246 | 0.181 |
| 217 | cgtA | 0.203 | 0.197 | 0.140 | 0.180 |
| 218 | dkgB | 0.203 | 0.197 | 0.140 | 0.180 |
| 219 | OG1RF_11718 | 0.145 | 0.284 | 0.105 | 0.178 |
| 220 | fabZ | 0.203 | 0.153 | 0.176 | 0.177 |
| 221 | pmi | 0.261 | 0.262 | 0.000 | 0.174 |
| 222 | gyrA2 | 0.145 | 0.131 | 0.246 | 0.174 |
| 223 | atpB | 0.145 | 0.197 |  | 0.171 |
| 224 | OG1RF_11397 | 0.174 | 0.197 | 0.140 | 0.170 |
| 225 | alsS | 0.203 | 0.197 | 0.105 | 0.168 |
| 226 | mtnN | 0.174 | 0.153 | 0.176 | 0.168 |
| 227 | pcp | 0.145 | 0.175 | 0.176 | 0.165 |
| 228 | rplV | 0.174 | 0.000 | 0.316 | 0.163 |
| 229 | hpt | 0.174 | 0.175 | 0.140 | 0.163 |
| 230 | murA | 0.174 | 0.175 | 0.140 | 0.163 |
| 231 | OG1RF_10839 | 0.203 | 0.175 | 0.105 | 0.161 |
| 232 | OG1RF_11999 | 0.203 | 0.175 | 0.105 | 0.161 |
| 233 | azoR | 0.174 | 0.131 | 0.176 | 0.160 |
| 234 | deoB | 0.174 | 0.131 | 0.176 | 0.160 |
| 235 | infC | 0.174 | 0.131 | 0.176 | 0.160 |
| 236 | rumA | 0.261 | 0.218 | 0.000 | 0.160 |
| 237 | aspS | 0.232 | 0.000 | 0.246 | 0.159 |
| 238 | pheS | 0.174 | 0.197 | 0.105 | 0.159 |
| 239 | purR | 0.174 | 0.197 | 0.105 | 0.159 |
| 240 | pflA | 0.145 | 0.153 | 0.176 | 0.158 |
| 241 | rex | 0.174 | 0.153 | 0.140 | 0.156 |
| 242 | OG1RF_10871 | 0.203 | 0.262 | 0.000 | 0.155 |
| 243 | fmt | 0.203 | 0.153 | 0.105 | 0.154 |
| 244 | yeeN | 0.203 | 0.153 | 0.105 | 0.154 |
| 245 | rpsS | 0.000 | 0.175 | 0.281 | 0.152 |
| 246 | yqeH | 0.174 | 0.175 | 0.105 | 0.151 |
| 247 | aspC2 | 0.145 | 0.306 | 0.000 | 0.150 |
| 248 | ddl | 0.145 | 0.197 | 0.105 | 0.149 |
| 249 | relA | 0.145 | 0.197 | 0.105 | 0.149 |
| 250 | oppD | 0.203 | 0.240 | 0.000 | 0.148 |
| 251 | OG1RF_12508 | 0.290 | 0.153 | 0.000 | 0.148 |
| 252 | OG1RF_12001 | 0.174 | 0.262 | 0.000 | 0.145 |
| 253 | aroB | 0.174 | 0.153 | 0.105 | 0.144 |
| 254 | fabZ | 0.174 | 0.153 | 0.105 | 0.144 |
| 255 | hslO | 0.174 | 0.153 | 0.105 | 0.144 |
| 256 | murA | 0.174 | 0.153 | 0.105 | 0.144 |
| 257 | dapE | 0.145 | 0.175 | 0.105 | 0.142 |
| 258 | gap | 0.145 | 0.175 | 0.105 | 0.142 |
| 259 | menB | 0.145 | 0.175 | 0.105 | 0.142 |
| 260 | OG1RF_12508 | 0.000 | 0.284 | 0.140 | 0.141 |
| 261 | OG1RF_10126 | 0.203 | 0.218 | 0.000 | 0.141 |
| 262 | tpiA | 0.000 | 0.175 | 0.246 | 0.140 |
| 263 | rfbD | 0.174 | 0.000 | 0.246 | 0.140 |
| 264 | fhs | 0.261 | 0.153 | 0.000 | 0.138 |
| 265 | nifJ | 0.261 | 0.153 | 0.000 | 0.138 |
| 266 | rpsR | 0.000 | 0.197 | 0.211 | 0.136 |
| 267 | OG1RF_11724 | 0.145 | 0.153 | 0.105 | 0.134 |
| 268 | glk | 0.290 | 0.000 | 0.105 | 0.132 |
| 269 | OG1RF_12441 | 0.174 | 0.218 | 0.000 | 0.131 |
| 270 | plsX | 0.000 | 0.175 | 0.211 | 0.128 |
| 271 | rpsF | 0.000 | 0.175 | 0.211 | 0.128 |
| 272 | pabC | 0.232 | 0.153 | 0.000 | 0.128 |
| 273 | parB | 0.232 | 0.153 | 0.000 | 0.128 |
| 274 | rplR | 0.174 | 0.000 | 0.211 | 0.128 |
| 275 | aroC | 0.145 | 0.131 | 0.105 | 0.127 |
| 276 | OG1RF_10897 | 0.145 | 0.131 | 0.105 | 0.127 |
| 277 | sufB | 0.232 | 0.000 | 0.140 | 0.124 |
| 278 | deoC | 0.174 | 0.197 | 0.000 | 0.124 |
| 279 | OG1RF_10191 | 0.174 | 0.197 | 0.000 | 0.124 |
| 280 | queA | 0.203 | 0.153 | 0.000 | 0.119 |
| 281 | rplU | 0.145 | 0.000 | 0.211 | 0.119 |
| 282 | rmlA | 0.000 | 0.175 | 0.176 | 0.117 |
| 283 | groS | 0.174 | 0.000 | 0.176 | 0.117 |
| 284 | aspC | 0.174 | 0.175 | 0.000 | 0.116 |
| 285 | cysS | 0.174 | 0.175 | 0.000 | 0.116 |
| 286 | thiI | 0.174 | 0.175 | 0.000 | 0.116 |
| 287 | atoB | 0.232 | 0.000 |  | 0.116 |
| 288 | ftsE | 0.145 | 0.197 | 0.000 | 0.114 |
| 289 | glyQ | 0.000 | 0.131 | 0.211 | 0.114 |
| 290 | pepQ2 | 0.000 | 0.153 | 0.176 | 0.109 |
| 291 | rplX | 0.000 | 0.153 | 0.176 | 0.109 |
| 292 | aroA | 0.174 | 0.153 | 0.000 | 0.109 |
| 293 | rnr | 0.174 | 0.153 | 0.000 | 0.109 |
| 294 | rplP | 0.145 | 0.000 | 0.176 | 0.107 |
| 295 | ftsY | 0.145 | 0.175 | 0.000 | 0.107 |
| 296 | pepC | 0.000 | 0.175 | 0.140 | 0.105 |
| 297 | rpsL | 0.174 | 0.000 | 0.140 | 0.105 |
| 298 | OG1RF_11332 | 0.203 | 0.000 | 0.105 | 0.103 |
| 299 | srpG2 | 0.203 | 0.000 | 0.105 | 0.103 |
| 300 | OG1RF_11029 | 0.174 | 0.131 | 0.000 | 0.102 |
| 301 | adh3 | 0.145 | 0.153 | 0.000 | 0.099 |
| 302 | gpsA | 0.145 | 0.153 | 0.000 | 0.099 |
| 303 | murG | 0.145 | 0.153 | 0.000 | 0.099 |
| 304 | nfo | 0.145 | 0.153 | 0.000 | 0.099 |
| 305 | ptsG | 0.145 | 0.153 | 0.000 | 0.099 |
| 306 | OG1RF_11908 | 0.000 | 0.153 | 0.140 | 0.098 |
| 307 | glxR | 0.000 | 0.175 | 0.105 | 0.093 |
| 308 | serS` | 0.000 | 0.175 | 0.105 | 0.093 |
| 309 | nhaC | 0.174 | 0.000 | 0.105 | 0.093 |
| 310 | OG1RF_11643 | 0.174 | 0.000 | 0.105 | 0.093 |
| 311 | panE2 | 0.174 | 0.000 | 0.105 | 0.093 |
| 312 | trxA | 0.174 | 0.000 | 0.105 | 0.093 |
| 313 | gshAB | 0.145 | 0.131 | 0.000 | 0.092 |
| 314 | murE | 0.145 | 0.131 | 0.000 | 0.092 |
| 315 | OG1RF_10780 | 0.145 | 0.131 | 0.000 | 0.092 |
| 316 | nagA | 0.000 | 0.153 | 0.105 | 0.086 |
| 317 | OG1RF_10929 | 0.000 | 0.153 | 0.105 | 0.086 |
| 318 | OG1RF_11749 | 0.145 | 0.000 | 0.105 | 0.083 |
| 319 | oppC | 0.145 | 0.000 | 0.105 | 0.083 |
| 320 | rbfA | 0.145 | 0.000 | 0.105 | 0.083 |
| 321 | OG1RF_10965 | 0.000 | 0.131 | 0.105 | 0.079 |
| 322 | luxS | 0.000 | 0.000 | 0.211 | 0.070 |
| 323 | rpsP | 0.000 | 0.000 | 0.211 | 0.070 |
| 324 | yfiA | 0.000 | 0.000 | 0.211 | 0.070 |
| 325 | OG1RF_11169 | 0.203 | 0.000 | 0.000 | 0.068 |
| 326 | parE | 0.203 | 0.000 | 0.000 | 0.068 |
| 327 | sun | 0.203 | 0.000 | 0.000 | 0.068 |
| 328 | ymdA | 0.000 | 0.197 | 0.000 | 0.066 |
| 329 | cspA | 0.000 | 0.000 | 0.176 | 0.059 |
| 330 | nadE2 | 0.000 | 0.000 | 0.176 | 0.059 |
| 331 | OG1RF_10206 | 0.000 | 0.000 | 0.176 | 0.059 |
| 332 | OG1RF_10477 | 0.000 | 0.000 | 0.176 | 0.059 |
| 333 | rplY | 0.000 | 0.000 | 0.176 | 0.059 |
| 334 | tig3 | 0.000 | 0.000 | 0.176 | 0.059 |
| 335 | trmFO | 0.000 | 0.000 | 0.176 | 0.059 |
| 336 | cmk | 0.000 | 0.175 | 0.000 | 0.058 |
| 337 | mreC | 0.000 | 0.175 | 0.000 | 0.058 |
| 338 | nusG | 0.000 | 0.175 | 0.000 | 0.058 |
| 339 | OG1RF_11715 | 0.000 | 0.175 | 0.000 | 0.058 |
| 340 | OG1RF_11855 | 0.000 | 0.175 | 0.000 | 0.058 |
| 341 | trcF | 0.000 | 0.175 | 0.000 | 0.058 |
| 342 | atpG | 0.174 | 0.000 | 0.000 | 0.058 |
| 343 | opuCC | 0.174 | 0.000 | 0.000 | 0.058 |
| 344 | codY | 0.000 | 0.153 | 0.000 | 0.051 |
| 345 | dapF | 0.000 | 0.153 | 0.000 | 0.051 |
| 346 | def | 0.000 | 0.153 | 0.000 | 0.051 |
| 347 | dnaA | 0.000 | 0.153 | 0.000 | 0.051 |
| 348 | era | 0.000 | 0.153 | 0.000 | 0.051 |
| 349 | fibA | 0.000 | 0.153 | 0.000 | 0.051 |
| 350 | gmk2 | 0.000 | 0.153 | 0.000 | 0.051 |
| 351 | OG1RF_10071 | 0.000 | 0.153 | 0.000 | 0.051 |
| 352 | rpiA | 0.000 | 0.153 | 0.000 | 0.051 |
| 353 | aspB2 | 0.145 | 0.000 | 0.000 | 0.048 |
| 354 | galE3 | 0.145 | 0.000 | 0.000 | 0.048 |
| 355 | nadE2 | 0.145 | 0.000 | 0.000 | 0.048 |
| 356 | OG1RF_10181 | 0.145 | 0.000 | 0.000 | 0.048 |
| 357 | OG1RF_11168 | 0.145 | 0.000 | 0.000 | 0.048 |
| 358 | OG1RF_11582 | 0.145 | 0.000 | 0.000 | 0.048 |
| 359 | OG1RF_11752 | 0.145 | 0.000 | 0.000 | 0.048 |
| 360 | OG1RF_11847 | 0.145 | 0.000 | 0.000 | 0.048 |
| 361 | OG1RF_11869 | 0.145 | 0.000 | 0.000 | 0.048 |
| 362 | OG1RF_12306 | 0.145 | 0.000 | 0.000 | 0.048 |
| 363 | pdp | 0.145 | 0.000 | 0.000 | 0.048 |
| 364 | fabK | 0.000 | 0.000 | 0.140 | 0.047 |
| 365 | mnmE | 0.000 | 0.000 | 0.140 | 0.047 |
| 366 | OG1RF_10922 | 0.000 | 0.000 | 0.140 | 0.047 |
| 367 | OG1RF_11289 | 0.000 | 0.000 | 0.140 | 0.047 |
| 368 | rpmC | 0.000 | 0.000 | 0.140 | 0.047 |
| 369 | rpsQ | 0.000 | 0.000 | 0.140 | 0.047 |
| 370 | murB | 0.000 | 0.131 | 0.000 | 0.044 |
| 371 | OG1RF_10337 | 0.000 | 0.131 | 0.000 | 0.044 |
| 372 | OG1RF_10952 | 0.000 | 0.131 | 0.000 | 0.044 |
| 373 | accA | 0.000 | 0.000 | 0.105 | 0.035 |
| 374 | asd | 0.000 | 0.000 | 0.105 | 0.035 |
| 375 | asp | 0.000 | 0.000 | 0.105 | 0.035 |
| 376 | atpF3 | 0.000 | 0.000 | 0.105 | 0.035 |
| 377 | ecfA | 0.000 | 0.000 | 0.105 | 0.035 |
| 378 | hprK | 0.000 | 0.000 | 0.105 | 0.035 |
| 379 | infA | 0.000 | 0.000 | 0.105 | 0.035 |
| 380 | manY | 0.000 | 0.000 | 0.105 | 0.035 |
| 381 | mscL | 0.000 | 0.000 | 0.105 | 0.035 |
| 382 | murC | 0.000 | 0.000 | 0.105 | 0.035 |
| 383 | OG1RF_10396 | 0.000 | 0.000 | 0.105 | 0.035 |
| 384 | OG1RF_11271 | 0.000 | 0.000 | 0.105 | 0.035 |
| 385 | OG1RF_11403 | 0.000 | 0.000 | 0.105 | 0.035 |
| 386 | OG1RF_11456 | 0.000 | 0.000 | 0.105 | 0.035 |
| 387 | OG1RF_12152 | 0.000 | 0.000 | 0.105 | 0.035 |
| 388 | rimP | 0.000 | 0.000 | 0.105 | 0.035 |
| 389 | rnpA | 0.000 | 0.000 | 0.105 | 0.035 |
| 390 | rpmD | 0.000 | 0.000 | 0.105 | 0.035 |
| 391 | rpsI3 | 0.000 | 0.000 | 0.105 | 0.035 |
| 392 | rpsT | 0.000 | 0.000 | 0.105 | 0.035 |
| 393 | ssb | 0.000 | 0.000 | 0.105 | 0.035 |
| 394 | tagD | 0.000 | 0.000 | 0.105 | 0.035 |
| 395 | trmB | 0.000 | 0.000 | 0.105 | 0.035 |
| 396 | trxB | 0.000 | 0.000 | 0.105 | 0.035 |
| 397 | ysxC | 0.000 | 0.000 | 0.105 | 0.035 |
