## Supplementary Table 5 for "The composition and function of *Enterococcus faecalis* membrane vesicles"

**Table S5**. **Proteins identified in membrane vesicle fraction by LC/MS/MS.** Proteins listed in descending order based on the abundance score. Abundance (A*) score is calculated as a number of unique peptides for each protein vs the number of total peptides within sample.

| A* position | protein name | A*1 | A*2 | A*3 | A*_avr_ |
| --- | --- | --- | --- | --- | --- |
| 1 | aad | **2.581** | **3.315** | **4.763** | **3.553** |
| 2 | groL | 3.125 | 2.957 | 4.102 | 3.394 |
| 3 | OG1RF_12384 | 1.427 | 2.867 | 3.837 | 2.710 |
| 4 | pbp1A | 1.495 | 2.778 | 3.440 | 2.571 |
| 5 | cnaB | 0.204 | 2.240 | 3.308 | 1.917 |
| 6 | OG1RF_10124 | 1.359 | 1.344 | 2.911 | 1.871 |
| 7 | OG1RF_12508 | 1.427 | 2.061 | 1.852 | 1.780 |
| 8 | OG1RF_12203 | 1.427 | 1.703 | 2.117 | 1.749 |
| 9 | OG1RF_10125 | 2.242 | 2.330 | 0.529 | 1.700 |
| 10 | penA | 1.223 | 1.703 | 2.117 | 1.681 |
| 11 | oppA | 1.223 | 1.971 | 1.720 | 1.638 |
| 12 | traC2 | 1.359 | 1.971 | 1.455 | 1.595 |
| 13 | OG1RF_12235 | 0.000 | 1.703 | 2.778 | 1.494 |
| 14 | pbp1B | 0.747 | 1.344 | 1.852 | 1.315 |
| 15 | prsA | 1.155 | 1.165 | 1.588 | 1.302 |
| 16 | rplB | 1.155 | 1.254 | 1.323 | 1.244 |
| 17 | tuf | 1.155 | 1.165 | 1.323 | 1.214 |
| 18 | eno | 0.000 | 1.613 | 1.985 | 1.199 |
| 19 | mreC | 0.611 | 1.254 | 1.455 | 1.107 |
| 20 | metQ | 0.747 | 0.986 | 1.455 | 1.063 |
| 21 | lepB | 0.747 | 0.896 | 1.455 | 1.033 |
| 22 | OG1RF_11679 | 0.883 | 1.075 | 1.058 | 1.006 |
| 23 | OG1RF_12509 | 1.291 | 1.613 | 0.000 | 0.968 |
| 24 | OG1RF_11718 | 0.611 | 1.075 | 1.191 | 0.959 |
| 25 | tyrS | 0.951 | 0.986 | 0.926 | 0.954 |
| 26 | OG1RF_11061 | 0.340 | 1.165 | 1.323 | 0.943 |
| 27 | fusA | 1.087 | 0.806 | 0.926 | 0.940 |
| 28 | ndh3 | 0.747 | 0.986 | 0.926 | 0.886 |
| 29 | OG1RF_11938 | 1.223 | 0.896 | 0.529 | 0.883 |
| 30 | psr | 0.543 | 0.806 | 1.191 | 0.847 |
| 31 | rpsB | 0.815 | 0.896 | 0.794 | 0.835 |
| 32 | lytR | 0.679 | 0.717 | 1.058 | 0.818 |
| 33 | pyk | 1.019 | 0.986 | 0.397 | 0.801 |
| 34 | OG1RF_12366 | 0.543 | 0.627 | 1.191 | 0.787 |
| 35 | OG1RF_11602 | 0.747 | 1.165 | 0.397 | 0.770 |
| 36 | gap2 | 0.747 | 0.627 | 0.926 | 0.767 |
| 37 | manX2 | 0.883 | 0.717 | 0.662 | 0.754 |
| 38 | OG1RF_10072 | 0.204 | 0.448 | 1.588 | 0.747 |
| 39 | pepA2 | 0.815 | 0.448 | 0.926 | 0.730 |
| 40 | dnaK | 0.747 | 0.627 | 0.794 | 0.723 |
| 41 | pbpC | 0.951 | 0.806 | 0.397 | 0.718 |
| 42 | pgk | 0.543 | 0.806 | 0.794 | 0.715 |
| 43 | OG1RF_12481 | 0.000 | 0.358 | 1.720 | 0.693 |
| 44 | OG1RF_11395 | 0.476 | 0.806 | 0.794 | 0.692 |
| 45 | rpoC | 0.951 | 0.538 | 0.529 | 0.673 |
| 46 | OG1RF_11907 | 0.679 | 0.538 | 0.794 | 0.670 |
| 47 | rpsC | 0.679 | 0.627 | 0.662 | 0.656 |
| 48 | OG1RF_11130 | 0.476 | 0.538 | 0.926 | 0.646 |
| 49 | fabI | 0.611 | 0.627 | 0.662 | 0.633 |
| 50 | pflB | 0.815 | 0.538 | 0.529 | 0.627 |
| 51 | lyzl6 | 0.272 | 0.806 | 0.794 | 0.624 |
| 52 | rpsG | 0.611 | 0.717 | 0.529 | 0.619 |
| 53 | atpA2 | 0.747 | 0.448 | 0.662 | 0.619 |
| 54 | OG1RF_10370 | 0.408 | 0.627 | 0.794 | 0.610 |
| 55 | rpsE | 0.611 | 0.627 | 0.529 | 0.589 |
| 56 | glnP | 1.019 | 0.627 | 0.000 | 0.549 |
| 57 | OG1RF_11354 | 0.340 | 0.627 | 0.662 | 0.543 |
| 58 | rpsD | 0.679 | 0.538 | 0.397 | 0.538 |
| 59 | oppD | 0.476 | 0.448 | 0.662 | 0.528 |
| 60 | pbp2A | 0.408 | 0.627 | 0.529 | 0.521 |
| 61 | OG1RF_10353 | 0.476 | 0.538 | 0.529 | 0.514 |
| 62 | pabC | 0.476 | 0.627 | 0.397 | 0.500 |
| 63 | atpD2 | 0.408 | 0.538 | 0.529 | 0.491 |
| 64 | rplT | 0.408 | 0.538 | 0.529 | 0.491 |
| 65 | divIVA2 | 0.340 | 0.448 | 0.662 | 0.483 |
| 66 | OG1RF_10021 | 0.340 | 0.448 | 0.662 | 0.483 |
| 67 | OG1RF_10897 | 0.408 | 0.448 | 0.529 | 0.462 |
| 68 | rplD | 0.476 | 0.358 | 0.529 | 0.454 |
| 69 | OG1RF_10136 | 0.815 | 0.000 | 0.529 | 0.448 |
| 70 | OG1RF_11807 | 0.272 | 0.538 | 0.529 | 0.446 |
| 71 | ftsH | 0.543 | 0.358 | 0.397 | 0.433 |
| 72 | rplS | 0.611 | 0.269 | 0.397 | 0.426 |
| 73 | OG1RF_11882 | 0.408 | 0.448 | 0.397 | 0.418 |
| 74 | rplF | 0.408 | 0.448 | 0.397 | 0.418 |
| 75 | mscL | 0.272 | 0.538 | 0.397 | 0.402 |
| 76 | htrA | 0.408 | 0.269 | 0.529 | 0.402 |
| 77 | rpsJ | 0.408 | 0.269 | 0.529 | 0.402 |
| 78 | pstS2 | 0.272 | 0.358 | 0.529 | 0.386 |
| 79 | secA | 0.883 | 0.269 | 0.000 | 0.384 |
| 80 | aceF | 0.611 | 0.538 | 0.000 | 0.383 |
| 81 | dps | 0.340 | 0.269 | 0.529 | 0.379 |
| 82 | glnA | 0.204 | 0.269 | 0.662 | 0.378 |
| 83 | OG1RF_12339 | 0.204 | 0.269 | 0.662 | 0.378 |
| 84 | OG1RF_12351 | 0.679 | 0.448 | 0.000 | 0.376 |
| 85 | pip | 0.204 | 0.896 | 0.000 | 0.367 |
| 86 | oppF | 0.340 | 0.358 | 0.397 | 0.365 |
| 87 | OG1RF_11240 | 0.408 | 0.269 | 0.397 | 0.358 |
| 88 | ldh | 0.272 | 0.269 | 0.529 | 0.357 |
| 89 | OG1RF_11255 | 0.272 | 0.358 | 0.397 | 0.342 |
| 90 | rplN | 0.272 | 0.358 | 0.397 | 0.342 |
| 91 | OG1RF_10126 | 0.747 | 0.269 | 0.000 | 0.339 |
| 92 | glmS | 0.340 | 0.269 | 0.397 | 0.335 |
| 93 | tig2 | 0.340 | 0.269 | 0.397 | 0.335 |
| 94 | hup | 0.204 | 0.269 | 0.529 | 0.334 |
| 95 | OG1RF_12464 | 0.000 | 0.448 | 0.529 | 0.326 |
| 96 | recA | 0.204 | 0.358 | 0.397 | 0.320 |
| 97 | rplL2 | 0.204 | 0.358 | 0.397 | 0.320 |
| 98 | srmB | 0.951 | 0.000 | 0.000 | 0.317 |
| 99 | rplV | 0.272 | 0.269 | 0.397 | 0.312 |
| 100 | OG1RF_12067 | 0.476 | 0.448 | 0.000 | 0.308 |
| 101 | OG1RF_11721 | 0.204 |  | 0.397 | 0.300 |
| 102 | OG1RF_10871 | 0.000 | 0.358 | 0.529 | 0.296 |
| 103 | rplU | 0.611 | 0.269 | 0.000 | 0.293 |
| 104 | ftsW2 | 0.204 | 0.269 | 0.397 | 0.290 |
| 105 | OG1RF_11697 | 0.204 | 0.269 | 0.397 | 0.290 |
| 106 | OG1RF_10486 | 0.204 | 0.000 | 0.662 | 0.288 |
| 107 | OG1RF_12540 | 0.476 | 0.358 | 0.000 | 0.278 |
| 108 | pfkA | 0.476 | 0.358 | 0.000 | 0.278 |
| 109 | celM | 0.543 | 0.269 | 0.000 | 0.271 |
| 110 | OG1RF_10350 | 0.272 | 0.538 | 0.000 | 0.270 |
| 111 | OG1RF_10479 | 0.408 | 0.000 | 0.397 | 0.268 |
| 112 | OG1RF_10869 | 0.000 | 0.269 | 0.529 | 0.266 |
| 113 | amiD | 0.340 | 0.448 | 0.000 | 0.263 |
| 114 | OG1RF_12031 | 0.340 | 0.448 | 0.000 | 0.263 |
| 115 | ptsI | 0.408 | 0.358 | 0.000 | 0.255 |
| 116 | rplC | 0.408 | 0.358 | 0.000 | 0.255 |
| 117 | tsf | 0.408 | 0.358 | 0.000 | 0.255 |
| 118 | OG1RF_12175 | 0.000 | 0.358 | 0.397 | 0.252 |
| 119 | OG1RF_10957 | 0.747 | 0.000 | 0.000 | 0.249 |
| 120 | OG1RF_12224 | 0.747 | 0.000 | 0.000 | 0.249 |
| 121 | rpoB | 0.747 | 0.000 | 0.000 | 0.249 |
| 122 | OG1RF_11751 | 0.272 | 0.448 | 0.000 | 0.240 |
| 123 | fruA | 0.340 | 0.358 | 0.000 | 0.233 |
| 124 | OG1RF_10191 | 0.340 | 0.358 | 0.000 | 0.233 |
| 125 | OG1RF_11455 | 0.340 | 0.358 | 0.000 | 0.233 |
| 126 | rplA | 0.408 | 0.269 | 0.000 | 0.225 |
| 127 | OG1RF_10867 | 0.000 | 0.269 | 0.397 | 0.222 |
| 128 | OG1RF_12417 | 0.000 | 0.269 | 0.397 | 0.222 |
| 129 | OG1RF_11723 | 0.272 | 0.358 | 0.000 | 0.210 |
| 130 | rplQ | 0.272 | 0.358 | 0.000 | 0.210 |
| 131 | ftsE | 0.340 | 0.269 | 0.000 | 0.203 |
| 132 | ndh2 | 0.340 | 0.269 | 0.000 | 0.203 |
| 133 | OG1RF_11094 | 0.340 | 0.269 | 0.000 | 0.203 |
| 134 | oppC | 0.340 | 0.269 | 0.000 | 0.203 |
| 135 | aroF | 0.204 | 0.000 | 0.397 | 0.200 |
| 136 | OG1RF_11059 | 0.204 | 0.000 | 0.397 | 0.200 |
| 137 | rplK | 0.204 | 0.000 | 0.397 | 0.200 |
| 138 | rplO | 0.204 | 0.000 | 0.397 | 0.200 |
| 139 | rseP | 0.204 | 0.000 | 0.397 | 0.200 |
| 140 | OG1RF_10218 | 0.204 | 0.358 | 0.000 | 0.187 |
| 141 | pdhB | 0.543 | 0.000 | 0.000 | 0.181 |
| 142 | divIVA | 0.272 | 0.269 | 0.000 | 0.180 |
| 143 | OG1RF_10495 | 0.272 | 0.269 | 0.000 | 0.180 |
| 144 | OG1RF_11390 | 0.272 | 0.269 | 0.000 | 0.180 |
| 145 | OG1RF_11999 | 0.272 | 0.269 | 0.000 | 0.180 |
| 146 | OG1RF_12091 | 0.272 | 0.269 | 0.000 | 0.180 |
| 147 | rpsM | 0.272 | 0.269 | 0.000 | 0.180 |
| 148 | OG1RF_12441 | 0.476 | 0.000 | 0.000 | 0.159 |
| 149 | opuAA | 0.476 | 0.000 | 0.000 | 0.159 |
| 150 | OG1RF_10839 | 0.204 | 0.269 | 0.000 | 0.158 |
| 151 | OG1RF_11464 | 0.204 | 0.269 | 0.000 | 0.158 |
| 152 | OG1RF_11698 | 0.204 | 0.269 | 0.000 | 0.158 |
| 153 | OG1RF_11710 | 0.204 | 0.269 | 0.000 | 0.158 |
| 154 | OG1RF_12152 | 0.204 | 0.269 | 0.000 | 0.158 |
| 155 | rpsH | 0.204 | 0.269 | 0.000 | 0.158 |
| 156 | rplE | 0.408 | 0.000 | 0.000 | 0.136 |
| 157 | fabZ | 0.408 | 0.000 | 0.000 | 0.136 |
| 158 | guaA | 0.408 | 0.000 | 0.000 | 0.136 |
| 159 | ndh | 0.408 | 0.000 | 0.000 | 0.136 |
| 160 | purA | 0.408 | 0.000 | 0.000 | 0.136 |
| 161 | pyrG | 0.408 | 0.000 | 0.000 | 0.136 |
| 162 | rplJ | 0.408 | 0.000 | 0.000 | 0.136 |
| 163 | rplW | 0.408 | 0.000 | 0.000 | 0.136 |
| 164 | rpsA | 0.408 | 0.000 | 0.000 | 0.136 |
| 165 | OG1RF_11056 | 0.000 | 0.358 | 0.000 | 0.119 |
| 166 | OG1RF_11062 | 0.000 | 0.358 | 0.000 | 0.119 |
| 167 | atpA | 0.340 | 0.000 | 0.000 | 0.113 |
| 168 | atpF3 | 0.340 | 0.000 | 0.000 | 0.113 |
| 169 | fbaB | 0.340 | 0.000 | 0.000 | 0.113 |
| 170 | infB | 0.340 | 0.000 | 0.000 | 0.113 |
| 171 | OG1RF_11761 | 0.340 | 0.000 | 0.000 | 0.113 |
| 172 | OG1RF_12392 | 0.340 | 0.000 | 0.000 | 0.113 |
| 173 | rpoA | 0.340 | 0.000 | 0.000 | 0.113 |
| 174 | tyrS | 0.340 | 0.000 | 0.000 | 0.113 |
| 175 | rplM | 0.272 | 0.000 | 0.000 | 0.091 |
| 176 | atpG | 0.272 | 0.000 | 0.000 | 0.091 |
| 177 | clpX | 0.272 | 0.000 | 0.000 | 0.091 |
| 178 | fabF | 0.272 | 0.000 | 0.000 | 0.091 |
| 179 | ftsA | 0.272 | 0.000 | 0.000 | 0.091 |
| 180 | ftsX | 0.272 | 0.000 | 0.000 | 0.091 |
| 181 | ftsZ | 0.272 | 0.000 | 0.000 | 0.091 |
| 182 | gatB | 0.272 | 0.000 | 0.000 | 0.091 |
| 183 | lpd | 0.272 | 0.000 | 0.000 | 0.091 |
| 184 | OG1RF_10127 | 0.272 | 0.000 | 0.000 | 0.091 |
| 185 | OG1RF_10977 | 0.272 | 0.000 | 0.000 | 0.091 |
| 186 | OG1RF_11506 | 0.272 | 0.000 | 0.000 | 0.091 |
| 187 | pgcA | 0.272 | 0.000 | 0.000 | 0.091 |
| 188 | pgi | 0.272 | 0.000 | 0.000 | 0.091 |
| 189 | nhaC | 0.000 | 0.269 | 0.000 | 0.090 |
| 190 | ftsQ | 0.000 | 0.269 | 0.000 | 0.090 |
| 191 | luxS | 0.000 | 0.269 | 0.000 | 0.090 |
| 192 | OG1RF_10281 | 0.000 | 0.269 | 0.000 | 0.090 |
| 193 | OG1RF_10285 | 0.000 | 0.269 | 0.000 | 0.090 |
| 194 | OG1RF_10540 | 0.000 | 0.269 | 0.000 | 0.090 |
| 195 | OG1RF_10875 | 0.000 | 0.269 | 0.000 | 0.090 |
| 196 | OG1RF_11034 | 0.000 | 0.269 | 0.000 | 0.090 |
| 197 | OG1RF_11060 | 0.000 | 0.269 | 0.000 | 0.090 |
| 198 | OG1RF_11722 | 0.000 | 0.269 | 0.000 | 0.090 |
| 199 | pstB2 | 0.000 | 0.269 | 0.000 | 0.090 |
| 200 | rpoD | 0.000 | 0.269 | 0.000 | 0.090 |
| 201 | ubiD | 0.000 | 0.269 | 0.000 | 0.090 |
| 202 | ziaA | 0.000 | 0.269 | 0.000 | 0.090 |
| 203 | OG1RF_10306 | 0.204 | 0.000 | 0.000 | 0.068 |
| 204 | OG1RF_10489 | 0.204 | 0.000 | 0.000 | 0.068 |
| 205 | OG1RF_11090 | 0.204 | 0.000 | 0.000 | 0.068 |
| 206 | gdhA | 0.204 | 0.000 | 0.000 | 0.068 |
| 207 | gpmA | 0.204 | 0.000 | 0.000 | 0.068 |
| 208 | OG1RF_10023 | 0.204 | 0.000 | 0.000 | 0.068 |
| 209 | OG1RF_10040 | 0.204 | 0.000 | 0.000 | 0.068 |
| 210 | OG1RF_10069 | 0.204 | 0.000 | 0.000 | 0.068 |
| 211 | OG1RF_10929 | 0.204 | 0.000 | 0.000 | 0.068 |
| 212 | OG1RF_11150 | 0.204 | 0.000 | 0.000 | 0.068 |
| 213 | OG1RF_11396 | 0.204 | 0.000 | 0.000 | 0.068 |
| 214 | OG1RF_12066 | 0.204 | 0.000 | 0.000 | 0.068 |
| 215 | OG1RF_12374 | 0.204 | 0.000 | 0.000 | 0.068 |
| 216 | pdhA | 0.204 | 0.000 | 0.000 | 0.068 |
| 217 | prs1 | 0.204 | 0.000 | 0.000 | 0.068 |
| 218 | pstB | 0.204 | 0.000 | 0.000 | 0.068 |
| 219 | pta | 0.204 | 0.000 | 0.000 | 0.068 |
| 220 | ptsG | 0.204 | 0.000 | 0.000 | 0.068 |
| 221 | pyrH | 0.204 | 0.000 | 0.000 | 0.068 |
| 222 | secY | 0.204 | 0.000 | 0.000 | 0.068 |
| 223 | srtA | 0.204 | 0.000 | 0.000 | 0.068 |
| 224 | sun | 0.204 | 0.000 | 0.000 | 0.068 |
| 225 | typA | 0.204 | 0.000 | 0.000 | 0.068 |
